# Endogenous recapitulation of Alzheimer’s disease neuropathology through human 3D direct neuronal reprogramming

**DOI:** 10.1101/2023.05.24.542155

**Authors:** Zhao Sun, Ji-Sun Kwon, Yudong Ren, Shawei Chen, Kitra Cates, Xinguo Lu, Courtney K. Walker, Hande Karahan, Sanja Sviben, James A J Fitzpatrick, Clarissa Valdez, Henry Houlden, Celeste M. Karch, Randall J. Bateman, Chihiro Sato, Steven J. Mennerick, Marc I. Diamond, Jungsu Kim, Rudolph E. Tanzi, David M. Holtzman, Andrew S. Yoo

## Abstract

Alzheimer’s disease (AD) is a neurodegenerative disorder that primarily affects elderly individuals, and is characterized by hallmark neuronal pathologies including extracellular amyloid-β (Aβ) plaque deposition, intracellular tau tangles, and neuronal death. However, recapitulating these age-associated neuronal pathologies in patient-derived neurons has remained a significant challenge, especially for late-onset AD (LOAD), the most common form of the disorder. Here, we applied the high efficiency microRNA-mediated direct neuronal reprogramming of fibroblasts from AD patients to generate cortical neurons in three-dimensional (3D) Matrigel and self-assembled neuronal spheroids. Our findings indicate that neurons and spheroids reprogrammed from both autosomal dominant AD (ADAD) and LOAD patients exhibited AD-like phenotypes linked to neurons, including extracellular Aβ deposition, dystrophic neurites with hyperphosphorylated, K63-ubiquitin-positive, seed-competent tau, and spontaneous neuronal death in culture. Moreover, treatment with β- or γ-secretase inhibitors in LOAD patient-derived neurons and spheroids before Aβ deposit formation significantly lowered Aβ deposition, as well as tauopathy and neurodegeneration. However, the same treatment after the cells already formed Aβ deposits only had a mild effect. Additionally, inhibiting the synthesis of age-associated retrotransposable elements (RTEs) by treating LOAD neurons and spheroids with the reverse transcriptase inhibitor, lamivudine, alleviated AD neuropathology. Overall, our results demonstrate that direct neuronal reprogramming of AD patient fibroblasts in a 3D environment can capture age-related neuropathology and reflect the interplay between Aβ accumulation, tau dysregulation, and neuronal death. Moreover, miRNA-based 3D neuronal conversion provides a human-relevant AD model that can be used to identify compounds that can potentially ameliorate AD-associated pathologies and neurodegeneration.

## Introduction

AD is the most common form of dementia and is accompanied by key features: extracellular deposition of Aβ plaques, formation of intracellular neurofibrillary tangles, and neuronal cell death^1,2^. The predominant approaches for modeling these AD pathological hallmarks have been transgenic cellular and animal models overexpressing ADAD-associated mutations^3–5^, leaving the LOAD relatively underexplored. LOAD accounts for over 95% of all AD cases and is a heterogeneous disease associated with a variety of risk factors, such as age, genetic predisposition, sex, stroke, and other factors^6, 7^. Among these, aging is the most significant risk factor, in which the risk increases exponentially with advanced age over 65^8, 9^. Due to the complex and heterogeneous nature of LOAD, patient-specific modelling using transgenic animals and cellular approaches has been challenging. Therefore, there is an urgent need to develop a patient-based model to study disease pathogenesis and understand its underlying mechanisms.

Recent advances in stem cell and reprogramming technologies have enabled the generation of various human cell types, including neurons differentiated from patient-specific induced pluripotent stem cells (iPSCs)^10^. Although these iPSC-derived neurons hold great promise for modelling and studying neurological development and diseases, induction of pluripotency in somatic cells erases the age signatures of the donor cells, resulting in iPSC-derived neurons reflecting embryonic identity^11^. This imposes challenges for modelling age-associated diseases such as LOAD, as it does not account for human age as the major contributing risk factor^12, 13^. For example, healthy adult human neurons express 4-repeat (4R) and 3-repeat (3R) tau isoforms at approximately 1:1 ratio^14, 15^. However, iPSC-derived neurons only express marginal levels of 4R tau isoforms, similar to fetal neurons^16, 17^. iPSC-derived neurons from AD patients exhibit increased levels of Aβ42, active GSK-3β, phosphorylated tau, endosomal abnormalities, and oxidative stress^18–20^, but no late-stage neuropathological features, such as neurofibrillary tau tangles and neurodegeneration. Therefore, establishing a patient-derived neuronal system that retains the age signature from the donor may be critical to recapitulate the key neuropathologic events in AD.

Direct cell lineage reprogramming converts cells of interest from one lineage to another through forced expression of genetic factors. By bypassing the pluripotent induction and stem cell stages, direct reprogramming offers experimental advantages to aging-related disorders, since age signatures such as the epigenetic clock, telomere length, and transcriptomic changes are propagated from the starting cells to the reprogrammed cells^21, 22^. Brain-enriched microRNAs, miR-9/9* and miR-124 (miR-9/9*-124), were identified as neurogenic reprogramming effectors that directly convert human fibroblasts into neurons^23^. MiR-9/9*-124 induce chromatin reconfiguration, allowing for sequential steps of fibroblast identity erasure, followed by neuronal program activation^24, 25^. This miRNAs-induced neuronal fate can synergize with additional transcription factors (TFs) to guide the conversion to disease-relevant neuronal subtypes with high efficiency^23, 24, 26, 27^ and is suitable for studying neuron-intrinsic properties of aging or age-dependent pathology^16, 28, 29^. Recent findings also demonstrated that miRNA-induced cortical neurons endogenously express all six tau isoforms analogous to adult human brains and are capable of forming insoluble tau in the presence of a tau mutation^16^, providing the foundation for modeling of tau-related pathologies.

In this study, we tested the feasibility of modeling the neuropathology of AD by generating AD patient-derived cortical neurons, a neuronal subtype susceptible to AD^6^, while addressing the central question of whether the age-maintained, patient-derived neurons would be sufficient to manifest key neuronal pathologies of AD. Leveraging the high efficiency miRNAs-mediated chromatin remodeling and reprogramming^16, 24, 25^, we developed a 3D Matrigel-based cell culture system for generating aged cortical neurons and self-organized neuronal spheroids to capture secreted Aβ and facilitate plaque formation. 3D-cultured neurons and spheroids reprogrammed from both ADAD and LOAD patients endogenously exhibited extracellular Aβ deposits, tau dysregulation, and neurodegeneration.

Moreover, inhibiting Aβ formation in LOAD neurons and spheroids reduces Aβ deposition, tauopathy and neuronal cell death, demonstrating that the toxic effect of Aβ accumulation can be captured by reprogrammed neurons and spheroids. By RNA-sequencing (RNA-seq), we identified neuroinflammation as a top enriched pathway in LOAD spheroids. Lastly, we validated that inhibition of transposable element synthesis alleviates AD pathologies in patient-derived neurons and spheroids. In summary, our results demonstrate the robustness of 3D culture-based direct neuronal reprogramming of patient fibroblasts to capture key neuropathological features of AD.

## Results

### Development of 3D direct neuronal reprogramming of AD fibroblasts into cortical neurons (CNs) and spheroids

Fibroblast samples from 10 AD patients (4 ADAD and 6 LOAD) and 16 age- and sex-matched healthy control (HC) donors (Table S1) were reprogrammed with neurogenic effectors miR-9/9*-124, and TFs NEUROD2 and MYT1L to drive the conversion towards cortical neurons^16, 23^. Consistent with previous reports^23^, the reprogrammed cells were electrically active in 2D culture, showing robust inward and outward currents and multiple action potentials, as measured by whole-cell recording (Fig. S1A). To limit the diffusion of secreted Aβ in conventional 2D culture and maximize Aβ deposition^3, 20^, we developed a Matrigel-based 3D-culture system consisting of a thin gel to allow for high resolution imaging of reprogrammed neurons and self-assembled neuronal spheroids. For thin gel culture, cells were dissociated at post-induction day (PID) 7 and resuspended in 15% Matrigel, followed by 2-3 weeks of reprogramming to generate 3D-cultured cortical neurons (3D-CNs) (Fig. 1A). During reprogramming, 3D-CNs underwent morphological changes from fibroblasts to neurons (Fig. 1B and S1B) and showed increased expression of long genes, a transcriptomic feature unique to neurons^25, 30^ (Fig. S1C). 3D-CNs also showed decreased expression of fibroblast genes and upregulation of neuronal genes during reprogramming (Fig. S1D). Immunostaining revealed that 3D-CNs derived from both HCs and AD patients expressed neuronal markers including NCAM1, MAP2, and SLC17A7 (also known as VGLUT1) (Fig. 1C and Fig. S1E).

**Fig. 1.**
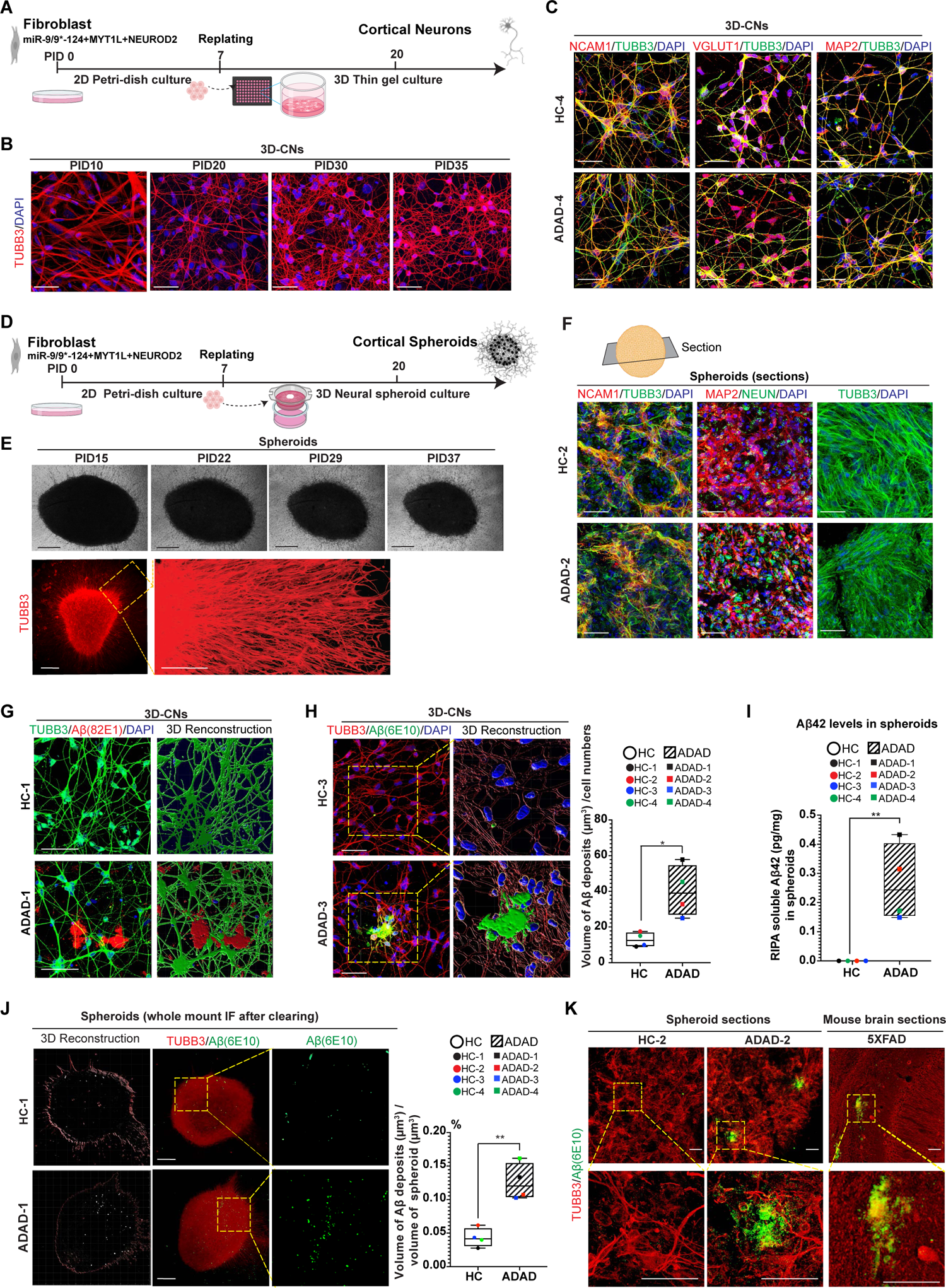
Direct reprogramming of AD patient fibroblasts into cortical neurons in 3D culture (3D-CNs) and neuronal spher-oids and ADAD 3D-CNs and spheroids showed higher amount of Aβ deposits. **A.** A schematic diagram of 3D-direct reprogramming of patient fibroblasts to cortical neurons in thin gel culture (PID: post-induction day). **B.** Representative immunofluorescence images with TUBB3 staining illustrating morphological changes during neuronal reprogramming in thin gel. Scale bar: 50 µm. **C.** Representative images of 3D-CNs stained with neuronal markers NCAM1, VGLUT1 and MAP2 at PID30 in thin gel. Scale bar: 50 µm. **D.** A schematic diagram of direct reprogramming of human fibroblasts to neuronal spheroids per transwell. **E.** Top: Representative phase contrast images of a neuronal spheroid at different PIDs of reprogramming. Scale bar: 500 µm. Bottom: A representative image of a neuronal spheroid at PID28 stained with TUBB3. Higher magnification view of the boxed region depicts neurite extensions radiating out from the core of the spheroids. Scale bar: 250 µm. **F.** Immunostaining in the sections of spheroids for neuronal markers NCAM1, MAP2 and NEUN and morphological marker TUBB3. Scale bar: 50 µm. **G.** Examples of extracellular Aβ deposits detected by 82E1 Aβ antibody in thin gel at PID30. Scale bar: 50 µm. **H.** Representative immunofluorescence images (left) and enlarged 3D reconstruction images (right) showing extracellular Aβ deposition (6E10) in thin gel at PID30. Quantification: total volume (µm^3^) of Aβ deposition measured using Imaris and was normalized to the total number of cells. Scale bar: 50 µm. **I.** Aβ42 levels were measured by electrochemiluminescence assay in ADAD and HC spheroids (PID28). **J.** Representative images of whole mount immunostaining of Aβ followed by clearing of ADAD spheroids and sex- and age-matched HC spheroids at PID28. 3D recon-struction was generated by Imaris and was used for quantification. Aβ deposition was quantified by measuring the total volume (µm^3^) of Aβ deposits labeled by 6E10 normalized to the volume (µm^3^) of spheroid labeled by TUBB3. Scale bar: 500 µm. **K.** Representative images of spheroid sections (PID28) and 5XFAD mouse brain cortex showing the extracellular Aβ deposition. Scale bar: 50 µm. For quantification in H, I, and J, n = 4 independent ADAD patients and 4 HC individuals. *p < 0.05 and **p < 0.01 were calculated by unpaired t-test.

For generating neuronal spheroids, cells were dissociated at PID7, pelleted, and transferred into a transwell insert to form one neuronal spheroid, followed by 2-3 weeks of culture in neuronal media (Fig. 1D). Reprogrammed spheroids exhibited extensive neurite outgrowth from the core of the spheroid (Fig. 1E). By 3-4 weeks, neuronal spheroids derived from HCs and AD patients displayed reduced expression of fibroblast marker genes and increased expression of long genes and neuronal markers compared to their starting fibroblasts, as assessed by RNA-seq (Fig.S1F and S1G) and confirmed by qPCR of neuronal markers such as *MAP2*, *MAPT* (also known as *TAU), SNAP25, ACTL6B* (also known as *BAF53B*, a marker of mature neurons^31^), *SLC17A6* (also known as *VGLUT1,* a glutamatergic neurons marker), *TBR1*(a cortical marker), and *POU3F2* (also known as *BRN2*, a cortical marker) (Fig. S1H). Immunofluorescence staining of PID28 spheroid sections revealed robust expression of NCAM1, MAP2 and NEUN in HC and AD spheroids (Fig. 1F). Collectively, these results confirm the neuronal identity of reprogrammed cells in 3D-direct reprogramming conditions.

### Aβ deposition in ADAD 3D-CNs and spheroids

To assess AD-like neuronal phenotypes, we performed immunostaining analyses of 3D-CNs derived from ADAD patient and age- and sex-matched HC individual using N-terminus specific Aβ antibody 82E1. Extracellular Aβ deposits with diameters of 10-50 µm were detected by 82E1 Aβ antibody in ADAD 3D-CNs at PID30 (Fig. 1G and Fig. S2A). We then examined Aβ deposition in the extracellular space of 3D-CNs from multiple ADAD patients with mutations in *PSEN1* or *APP* (Table S1) and age- and sex-matched HC fibroblast lines using another anti-Aβ antibody 6E10. Similar to the result of using 82E1 antibody, the extracellular deposition of Aβ in ADAD 3D-CNs was also detected by 6E10 Aβ antibody, with a threefold increase of extracellular Aβ deposit volume compared to HC 3D-CNs (Fig. 1H). To examine the Aβ deposition in spheroids, we first measured the Aβ42 level through electrochemiluminescence assay in spheroids after 4 weeks of reprogramming. In contrast to low Aβ42 levels detected in HC spheroids, ADAD spheroids showed significantly higher levels of Aβ42 (Fig. 1I), consistent with the impact of ADAD mutations on APP processing. Next, we performed whole mount immunostaining using Aβ antibody (6E10) on PID22 and PID28 spheroids followed by clearing of the spheroids to allow 3D fluorescent imaging. At both PID22 and PID28, ADAD spheroids exhibited a significantly higher amount of Aβ deposition compared to HC spheroids (Fig. 1J and S2B). In contrast to HC spheroids, however, ADAD spheroids showed a significant increase of Aβ deposition from PID22 to PID 28 (Fig. S2B). To confirm the extracellular deposition of Aβ in spheroids, cryo-sections of PID28 spheroids were immunostained using two different Aβ antibodies (6E10 and 82E1). Extracellular Aβ deposits were detected in ADAD spheroids by 6E10 or 82E1 antibodies, as observed in 5XFAD mouse brain sections (Fig. 1K and Fig. S2C). Transmission electron microscopy (TEM) with immunogold labeling against Aβ also showed gold signals in the extracellular domain in ADAD spheroids (Fig. S2E). Finally, to confirm that Aβ deposits observed in ADAD spheroids were generated through APP cleavage, we treated the spheroids with β-secretase inhibitor IV or DAPT from PID16 to 28, which led to the reduction of Aβ deposition (Fig. S2D). Taken together, these data demonstrate that ADAD 3D-CNs and spheroids contain elevated levels of extracellular Aβ deposition.

### Tau dysregulation in ADAD 3D-CNs and spheroids

In AD, tau protein becomes hyperphosphorylated, resulting in tau dissociating from microtubules and forming insoluble tau aggregates in the neurites and cell bodies^3,32^. To investigate the level of phosphorylated tau (p-tau) in ADAD 3D-CNs compared to age-matched HC 3D-CNs, we carried out immunostaining analyses using AT8 (phosphor-Ser202/Thr205) and PHF1 (phosphor-Ser396/Ser404) p-tau antibodies. Both ADAD and HC neurites exhibited p-tau signals similar to the tau islands patterns seen in primary neurons^16, 33, 34^ (Fig. 2A and Fig. S3A). However, ADAD 3D-CNs displayed significantly higher levels of p-tau compared to HC 3D-CNs (Fig. 2A). ADAD 3D-CNs also showed higher signals for MC1 antibody, which detects conformationally misfolded tau^35^, compared to HC 3D-CNs (Fig. 2B). In human AD brains, K63-linked ubiquitinated tau accumulates at high levels and is associated with enhanced tau seeding and propagation^36^. By performing immunostaining using K63-linked ubiquitin and p-tau (PHF1) antibodies, we found that ADAD 3D-CNs showed significantly higher volumes of K63-ubiquitin in PHF1-positive neurites compared to HC 3D-CNs (Fig. 2C).

**Fig. 2.**
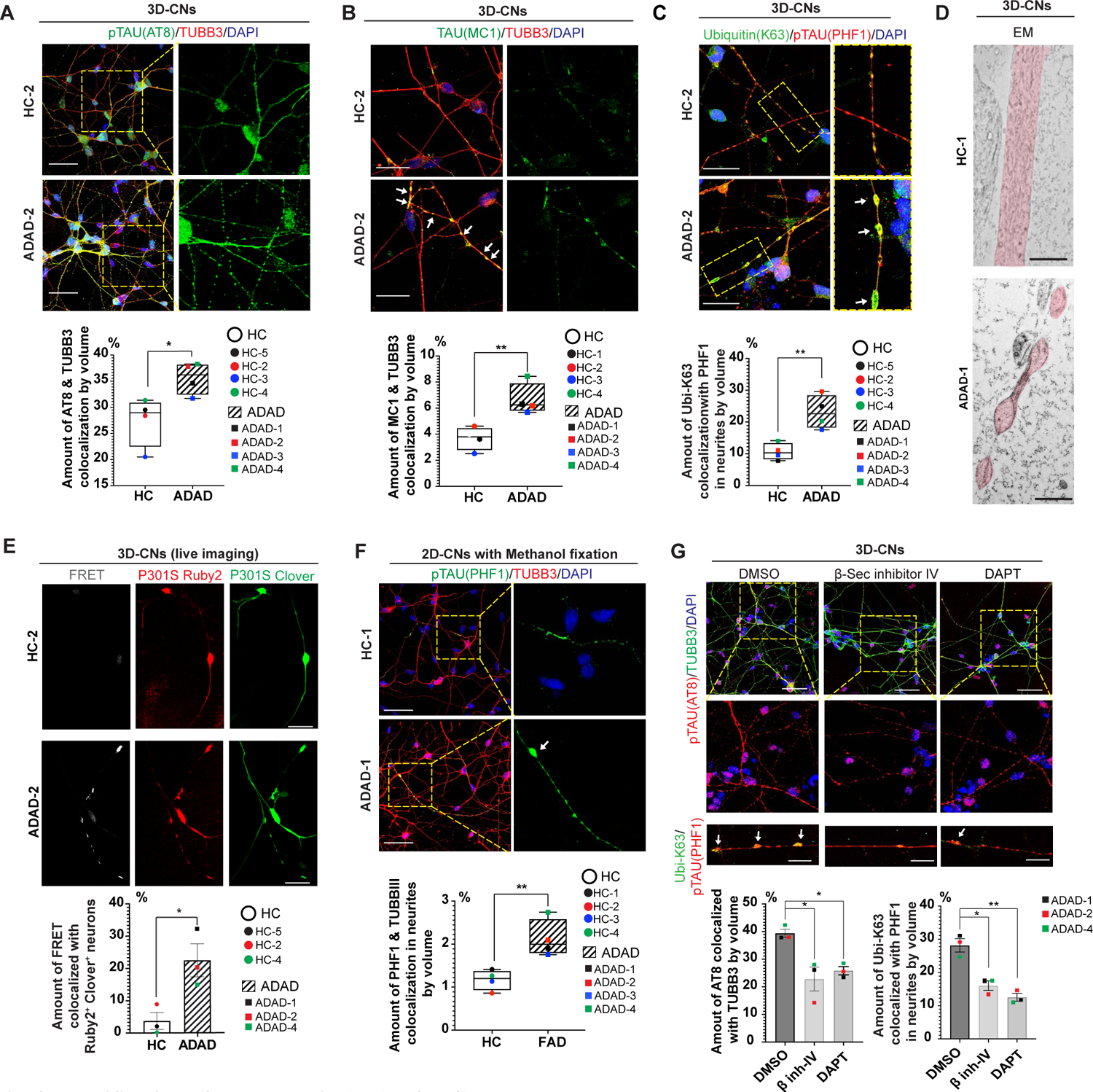
Identification of tauopathy in ADAD 3D-CNs. **A.** Immunofluorescence staining of p-tau (AT8 antibody) in ADAD and HC 3D-CNs (PID30). Scale bar: 50 µm. **B.** Misfolded tau was detected by immunostaining with MC1 tau antibody in ADAD and HC 3D-CNs at PID30. Arrows highlight the tau islands that are positive for MC1 antibody. Scale bar: 25 µm. **C.** Immunofluorescence images of pathogenic tau by co-staining of K63-linked ubiquitin (green) and p-tau (PHF1) (red) in ADAD and HC 3D-CNs. Arrows point to the enlarged beaded neurite bulges with PHF1 signal colocalized with K63-linked Ubiquitin. Scale bar: 25 µm. **D.** Representative transmission EM images of healthy neurites (top) and dystrophic “beaded” neurites (bottom) from HC and ADAD 3D-CNs, respectively. Scale bar: 800 nm. **E.** Representative live-cell FRET images for detecting seed-competent tau from HC and ADAD 3D-CNs at PID26. For each group (HC vs ADAD), the representative images were taken from same field with different channels. Scale bar: 25 µm. Data was shown by Mean ± SEM. **F.** Immunofluorescence images of p-tau (PHF1) in methanol-fixed 2D cultured cells revealing insoluble tau. Arrow points to the dystrophic tau bead. Scale bar: 25 µm. **G.** Immunofluorescence of p-tau (AT8) (top) and p-tau^+^ (PHF1) and K63-ubiquitin^+^ dystrophic neurites (bottom) in ADAD 3D-CNs treated with β-secretase inhibitor IV or DAPT. Arrows depict swelled tau blebs in 3D-CNs. Scale bar: 25 µm. For quantification in A, B, C and F, n = 4 independent ADAD patients and 4 HC individuals. For E and G, n = 3 ADAD patients and 3 HC individuals. Imaris was used to calculate the volumes marked by different antibodies. The amounts of TUBB3 that colocalized with AT8 p-tau (A and G top), MC1 tau (B) or insoluble PHF1 p-tau (F) (µm^3^/µm^3^), amounts of PHF1 p-tau that colocalized with K63-linked ubiquitin (C and G bottom) (µm^3^/µm^3^), and the amounts of FRET^+^ neurons as a fraction of Ruby2^+^ Clover^+^ neurons (E) (µm^3^/µm^3^) were calculated for statistics. *p < 0.05 and **p < 0.01 were calculated by unpaired t-test in A, B, C, E and F. Adjusted p-values were calculated by one-way ANOVA with Dunnett’s multiple comparisons test in G. * adjusted p < 0.05 and ** adjusted p < 0.01.

Importantly, “beading” or “blebbing” in neurites is a morphological sign of dystrophic neurites in AD brain^3,37, 38^. Notably, we detected spherical, beaded neurites in ADAD 3D-CNs and spheroids that are positive for p-tau/K63-ubiquitin staining (Fig. 2C and S3B, see by arrows). We further confirmed the presence of beaded dystrophic neurites in ADAD neurons by high-resolution TEM (Fig. 2D).

To test whether ADAD 3D-CNs would contain seed-competent tau that facilitates spreading of pathogenic tau, we carried out a Fluorescence Resonance Energy Transfer (FRET) assay by transducing tau FRET reporters composed of tau P301S-Ruby2 and -Clover reporters, which dimerize and transmit FRET in the presence of seed-competent tau species^39, 40^. As shown in Fig. 2E, ADAD 3D-CNs exhibited significantly increased FRET signals compared to HCs, suggesting that ADAD 3D-CNs contain seed-competent tau. Intriguingly, the FRET signal in ADAD 3D-CNs was specifically located in beaded dystrophic neurite regions (Fig. 2E). Methanol fixation has previously been used to remove soluble proteins, including tau^16, 41^. Immunostaining using PHF1 p-tau antibody in methanol fixed CNs indicated that ADAD CNs exhibited elevated amounts of insoluble p-tau (Fig. 2F).

Moreover, we also used GT-38 tau antibody which selectively labels the AD-specific tau strain, and its abundance is correlated with disease stage^42^. Immunostaining of ADAD 3D-CNs showed GT-38-labeled tau albeit with low occurrences (Fig. S3C). Finally, we tested whether Aβ deposits would underlie tau dysregulation in patient-derived neurons^3,6^ by treating ADAD 3D-CNs with β-secretase inhibitor IV or γ-secretase inhibitor (DAPT) to prevent APP processing. We found that inhibiting APP led to significant reduction of p-tau (AT8) and K63-ubiquitin^+^ p-tau (PHF1) levels in ADAD 3D-CNs (Fig. 2G). Altogether, our results indicate that ADAD neurons manifest tau pathology that is dependent on the formation of Aβ deposits.

### ADAD 3D-CNs and spheroids undergo spontaneous neurodegeneration

AD is a neurodegenerative disorder characterized by the loss of neurons, particularly in the hippocampus and cerebral cortex^43, 44^. To test if directly reprogrammed ADAD 3D-CNs would undergo spontaneous neurodegeneration, we examined cell death in 3D-CNs at multiple time points during reprogramming by Sytox-Green assay, a general cell death indicator. While both ADAD and HC reprogramming cells showed low levels of cell death prior to PID20, significantly higher levels of cell death were detected in ADAD 3D-CNs at PID30 and PID35, compared to HCs at same PIDs (Fig. 3A). These time points correspond to when neuronal identity is established during the conversion process (Fig. 1B and Fig. S1C). We also examined neuronal death in spheroids by carrying out TUNEL staining on sections of spheroids at PID28. Consistent with 3D-CNs, ADAD spheroids showed significantly higher cell death levels compared to HC spheroids (Fig. 3B). We further tested the specificity of the neuronal death phenotype by culturing ADAD spheroids (labeled by turboRFP) immediately adjacent to HC spheroids (labeled by eGFP). Remarkably, ADAD spheroids demonstrated stark shrinkage in size by 50% from PID17 to PID33, while HC spheroids exhibited only a 10% reduction during the same time interval (Fig. 3C, right). In contrast, two adjacent HC spheroids derived from independent control donors did not undergo significant shrinkage from PID17 to PID33 (Fig. 3C, left) indicating that AD-associated neurodegeneration was specifically recapitulated in ADAD spheroids. Since AD is also characterized by progressive degeneration of neurites^45^, we examined neurite outgrowth in ADAD and HC spheroids at PID22 and PID28. HC spheroids maintained consistent neurite length from PID22 to PID28, while ADAD spheroids showed significant retraction of neurite outgrowth at PID28 (Fig. 3D). Lastly, we tested whether the neuronal death would be a phenotype downstream of Aβ deposits. Treating ADAD spheroids with β- or γ-secretase inhibitors led to significant reduction of neuronal death (Fig 3E). Altogether, these results demonstrate that ADAD 3D-CNs and spheroids undergo spontaneous, Aβ-dependent neurodegeneration.

**Fig. 3.**
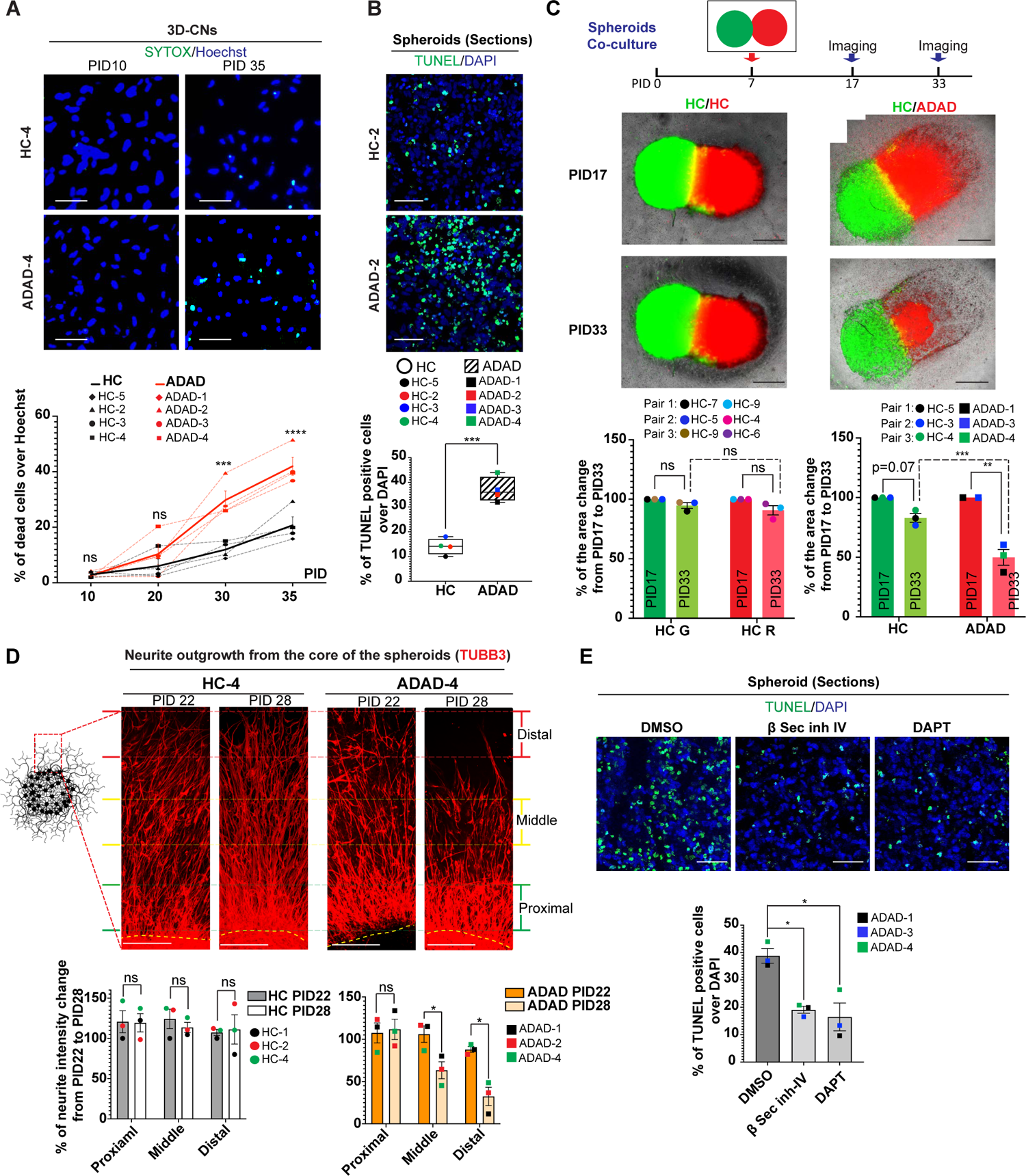
Severe neurodegeneration in ADAD 3D-CNs and spheroids. **A.** Representative immunofluorescence images of Sytox-positive cells in ADAD and HC 3D-CNs at different PIDs. Quantification shows the percentage of Sytox-positive cells over the number of Hoechst-positive cells. n = 4 ADAD patients and 4 HC individuals. Solid lines show the average of each group while dotted lines represent each individual. Mean ± SEM. p-values were calculated by two-way ANOVA with Šídák’s multiple comparisons test. ns = adjusted p > 0.05, *** adjusted p < 0.001 and **** adjusted p < 0.0001. Scale bar: 50 µm. **B.** Representative images of TUNEL staining in ADAD or HC spheroid sections. Quantifications show the percentage of TUNEL positive cells over DAPI. n = 4 ADAD patients and 4 HC individuals. ***p < 0.001 by unpaired t-test. Scale bar: 50 µm. **C.** Two spheroids were labeled by RFP or GFP and co-cultured starting at PID7. At PID17 and PID33, multiple live images were taken and stitched to display the whole spheroids. Left: Two spheroids derived from two different HC individuals are labeled by RFP or GFP. Right: ADAD spheroids were labeled by RFP and HC spheroids were labeled with GFP. The areas (µm^2^) of RFP-labeled and GFP-labeled spheroids at PID17 and PID33 were measured by Imaris. Left (HC/HC co-culture): n = 3 pairs (total 5 HC lines). Right (HC/ADAD co-culture): n = 3 pairs (3 ADAD patients and 3 HC individuals). Mean ± SEM. Adjusted-p values were calculated by two-way ANOVA with Šídák’s multiple comparisons test. ns = adjusted p > 0.05; **adjusted p < 0.01, and ***adjusted p < 0.001. HC G: HC spheroids with GFP; HC R: HC spheroids with RFP. Scale bar: 1 mm. **D.** Left: Cartoon illustration of the typical spheroids with neurite outgrowth from the core. Top: Representative TUBB3 immunostaining images show neurite outgrowth from HC and ADAD spheroids at PID22 and PID28. Yellow dashed lines indicate the border between the core and the neurites. Scale bar: 250 µm. Bottom: Quantifications show the percentage of intensity in proximal, middle, and distal regions of HC and ADAD neurites normalized by the intensity in the same region of PID22 HC-1 neurites (set as 100%). n = 3 ADAD patients and 3 HC individuals. Mean ± SEM. *p < 0.05 was calculated by multiple paired t-tests. **E.** Cell death in ADAD spheroids (PID28) was reduced by treating spheroids with β-secretase inhibitor IV or DAPT starting at PID 16. n = 3 ADAD patients. Mean ± SEM. *Adjusted p < 0.05 was calculated by one-way ANOVA with Dunnett’s multiple comparisons test. Scale bar: 50 µm.

### LOAD 3D-CNs and spheroids display increased Aβ deposition, tau dysregulation, dystrophic neurites, and neuronal cell death phenotypes

While reprogrammed ADAD neurons capture the adult-onset neuropathology resulting from disease-causing genetic mutations, it is not clear if LOAD samples would also manifest AD-related neuropathology. As such, we asked if the 3D neuronal reprogramming could be applied to LOAD samples to model AD-associated neurodegeneration. It is noteworthy that Aβ deposition is not limited to AD patients but can be also present in non-demented elderly individuals^46^. Thus, we first examined Aβ deposition in 3D-CNs from healthy individuals at different ages. While Aβ deposits were barely detected in the 3D-CNs derived from a young adult (22 years of age), Aβ deposition became more detectable with increasing ages (Fig S4A). Therefore, for comparing Aβ between LOAD and HC CNs, it is critical to match the donor ages. Both LOAD and HC 3D-CNs displayed extracellular Aβ deposits detected by 6E10 and 82E1 Aβ antibodies (Fig. 4A and Fig. S4B); however, LOAD 3D-CNs (6 independent patients, 60-89 years old) showed higher levels of Aβ deposition compared to HC 3D-CNs from six independent controls of similar age range (Fig. 4A). Similarly, LOAD spheroids from six independent patients exhibited significantly higher levels of extracellular Aβ deposits at PID28 compared to age-matched HC spheroids from six independent control individuals (Fig. 4B and Fig. S4C). These data demonstrate the feasibility of recapitulating Aβ deposition in LOAD and aged HC neurons via 3D neuronal reprogramming.

**Fig. 4.**
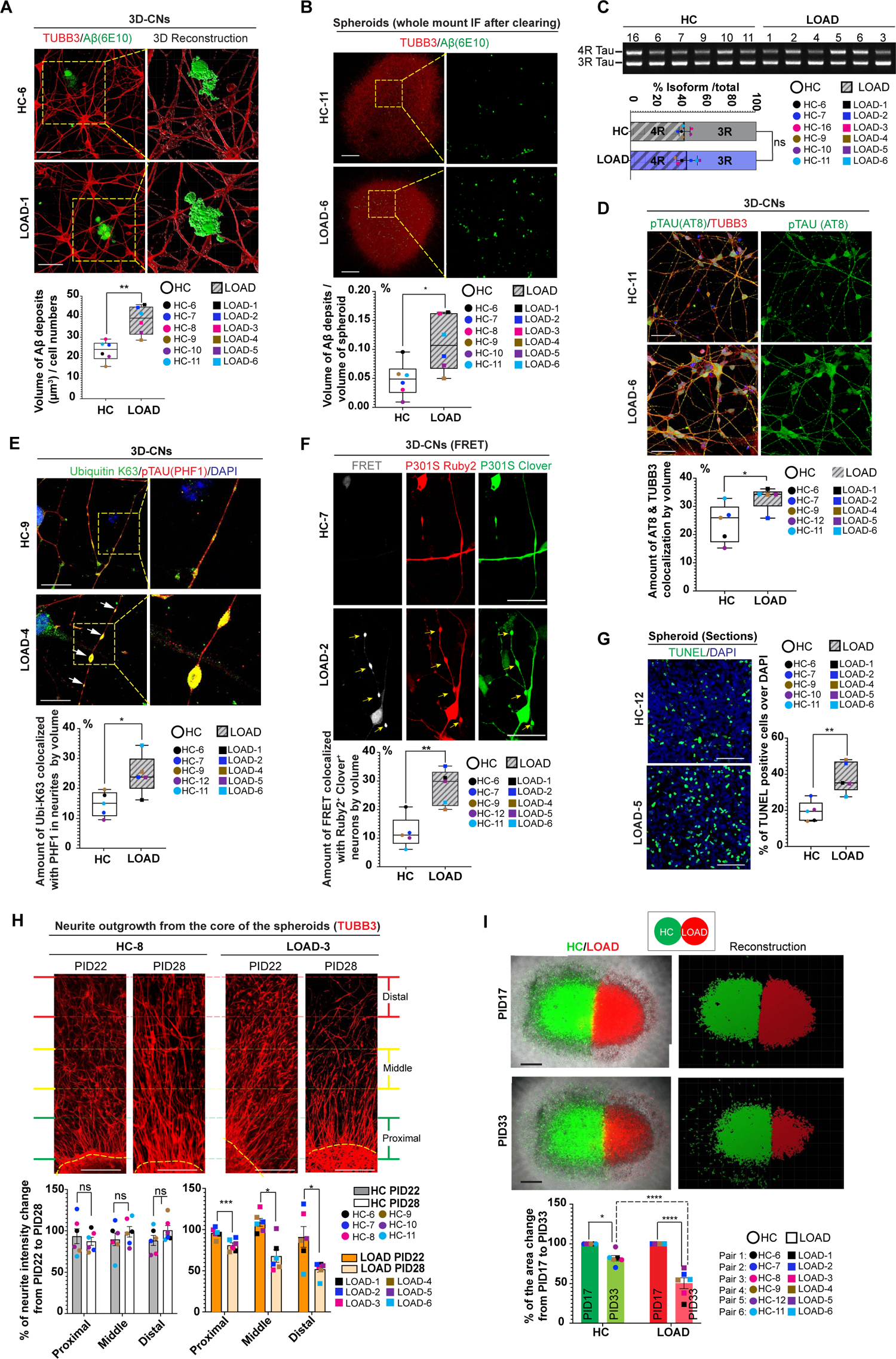
Elevated Aβ deposition, pathogenic tau, and neurodegeneration in LOAD 3D-CNs and spheroids. **A.** Representative immunofluorescence images of Aβ deposits in thin gel culture of PID30 LOAD and HC neurons. Boxed regions were highlighted on the right by 3D reconstruction showing extracellular Aβ deposition. Quantification: total volume (µm^3^) of Aβ deposits normalized to the total number of cells. Scale bar: 50 µm. **B.** Whole mount immunostaining with Aβ antibody (6E10) in PID28 LOAD and HC spheroids showed enhanced Aβ deposition in LOAD spheroids. Boxed regions were magnified at the right panels to show Aβ deposition. Volumes (µm^3^) of total Aβ deposits were normalized to the volumes (µm^3^) of TUBB3^+^ spheroid for quantification. Scale bar: 500 µm. **C.** PCR analysis of 3R and 4R tau isoforms in LOAD and HC 3D-CNs at PID25. Data was shown as Mean ± SEM. **D.** At PID30, LOAD 3D-CNs contained significantly more AT8^+^ p-tau than HC 3D-CNs. Scale bar: 50 µm. **E.** Co-staining of K63-specific ubiquitin (green) and p-tau (PHF1) (red) in the neurites (PID30 3D-CNs). Arrows highlight the swelled dystrophic neurite bulges containing K63-ubiquitin and tau signal in LOAD neurons. Scale bar: 25 µm. **F.** Live-cell imaging of FRET signal from HC and LOAD 3D-CNs containing Ruby2 and Clover reporters at PID28, and quantification of FRET signals. Scale bar: 25 µm. **G.** Representative images of TUNEL staining on sections of LOAD and HC spheroids (PID28). Scale bar: 50 µm. **H.** The neurite outgrowth at proximal, middle, and distal regions in LOAD and HC spheroids examined by immunofluorescence with TUBB3 antibody. TUBB3 signals in each region of the neurites were compared to same region in HC-6 spheroid at PID22 (set as 100%). Mean ± SEM. Scale bar: 250 µm. **I.** LOAD spheroids (labeled with RFP) and HC spheroids (labeled with GFP) were co-cultured at PID7 and fluorescence images were taken at PID17 and PID33. Spheroid size (area, µm^2^) at PID33 was normalized to its size at PID17. Scale bar: 1 mm. Mean ± SEM. For quantifications: n = 6 LOAD patients and 6 HC individuals in A, B, C, H and I, n = 5 LOAD patients and 5 HC individuals in D, E, F and G. Unpaired t-tests were used for calculating p-values in A, B, C, D, E, F and G (*p < 0.05, **p < 0.01 and ns p > 0.05). Multiple paired t-test was calculated for H (p < 0.05, ***p < 0.001 and ns p > 0.05). Two-way ANOVA with Šídák’s multiple comparisons test was used for I (*adjusted p < 0.05 and **** adjusted p < 0.0001).

In AD, tau tangles contain a mixture of 3R- and 4R-tau isoforms to adopt an AD-characteristic topology^14, 15^. Thus, endogenous expression of both 3R and 4R tau isoforms is a critical criterion when using reprogrammed human neurons for modeling tau pathology in AD. To examine if neurons reprogrammed from LOAD patients express both 3R and 4R tau isoforms, we carried out semi-quantitative PCR^16^. Both 3R- and 4R-tau isoforms were detected with no overt difference in the 3R to 4R ratio between LOAD and HC 3D-CNs (Fig. 4C). Several lines of evidence suggest that increased tau phosphorylation is a reflection of brain aging in general^47, 48^. Interestingly, we noted that the trend of increased tau phosphorylation is correlated with the increasing ages of control donors (Fig. S5A).

Phosphorylated tau (AT8) was detected in both LOAD 3D-CNs and aged-matched HC 3D-CNs when reprogrammed cells gained the neuronal identity at PID30 and PID35, but not at PID10 and 20 (Fig. S5B). Importantly in comparison between LOAD and HC, we found that LOAD 3D-CNs exhibited significantly higher p-tau levels compared to HC 3D-CNs, as assessed by immunostaining with AT8 and PHF1 antibodies (Fig. 4D and Fig. S5C). LOAD 3D-CNs also showed significantly higher signals for MC1 tau antibody which recognizes conformationally misfolded tau^35^ (Fig. S5D). Additionally, in LOAD 3D-CNs there was significant elevation of K63-ubiquitin^+^ pathogenic tau signals, primarily located in beaded neurites, compared to HC 3D-CNs (Fig 4E). Similar to ADAD 3D-CNs (Fig. 2E), we also detected seed-competent tau in LOAD 3D-CNs as measured by FRET signals, especially in the beaded neurite regions (Fig. 4F). Altogether, these results indicate that LOAD patient-derived neurons are capable of capturing tau dysregulation through 3D reprogramming.

Brain atrophy due to neuronal loss is a pathologic feature in LOAD patients^9^. While LOAD and HC reprogramming cells did not display apparent cell death during the early phase of reprogramming (PID20), both PID30 and 35 time points showed significantly higher levels of Sytox signals in LOAD 3D-CNs compared to HC 3D-CNs (Fig. S6A). Live cell tracking over time confirmed the severe neuronal loss in LOAD 3D-CNs compared with HC 3D-CNs at later stages of reprogramming when the neuronal identity was evoked (Fig. S6B). LOAD spheroids also presented a significant increase of cell death, as revealed by TUNEL staining (Fig. 4G), and disintegration of neurites over time (Fig. 4H), in comparison to HC spheroids. Finally, culturing RFP-labeled LOAD spheroids adjacent to GFP-labeled HC spheroids revealed a ∼ 50% reduction of the size of LOAD spheroids from PID17 to PID33, while only ∼15-20% reduction of size was observed in the HC counterparts over the same period (Fig. 4I). Taken together, our data demonstrate that LOAD 3D-CNs and spheroids recapitulate AD-associated neuropathology characterized by increased Aβ deposition, tau dysregulation, and neuronal death compared to age- and sex-matched HCs.

### Effect of inhibiting APP processing in LOAD 3D-CNs and spheroids on AD neuropathology

LOAD 3D-CNs and spheroids represent a previously unavailable patient neuron-based system that endogenously captures AD-associated neuropathology. We thus tested the interplay between Aβ deposition, tauopathy and neurodegeneration by examining effects of reducing Aβ deposition on tau and neurodegeneration. This was assessed at two time points of reprogramming. First, we started treating reprogramming cells with β- or γ-secretase inhibitors at PID16, a time point before the observed onset of Aβ deposition (Fig S4D). Treating LOAD spheroids with β- or γ-secretase inhibitors before the formation of Aβ deposits significantly reduced Aβ deposits in the spheroids by 80-90% and cleaved Aβ species in the media compared to DMSO (Fig. 5A and Fig. S4E). Second, we started to treat reprogramming cells with β- or γ-secretase inhibitors at PID22, when moderate amount of Aβ deposits is already present (Fig S4D). Inhibiting APP processing after Aβ deposition has already begun resulted in either a non-significant effect (by β-secretase inhibitor IV) or only mild reduction (∼30% by DAPT) on Aβ deposition (Fig. 5A). Next, we found that inhibition of Aβ formation before accumulation of Aβ deposits significantly decreased p-tau level and K63-ubiquitin/p-tau colocalization, whereas inhibition of Aβ formation at the later time point resulted in only non-significant effects on tau dysregulation (Fig. 5B). Lastly, we examined neuronal death in spheroids treated with β- or γ-secretase inhibitors. Similar to the effects on tau pathology, neuronal death in LOAD spheroids was reduced by nearly 50% when APP processing was inhibited starting at an early phase of reprogramming (PID16). However, secretase inhibitors had only minor effects on neuronal death if applied after the onset of Aβ aggregation (Fig. 5C). Together, these results demonstrate that the timing of blocking Aβ production is a crucial factor for the inhibition of Aβ formation and subsequent tauopathy and neuronal death in the LOAD 3D-CN and spheroid models.

**Fig. 5.**
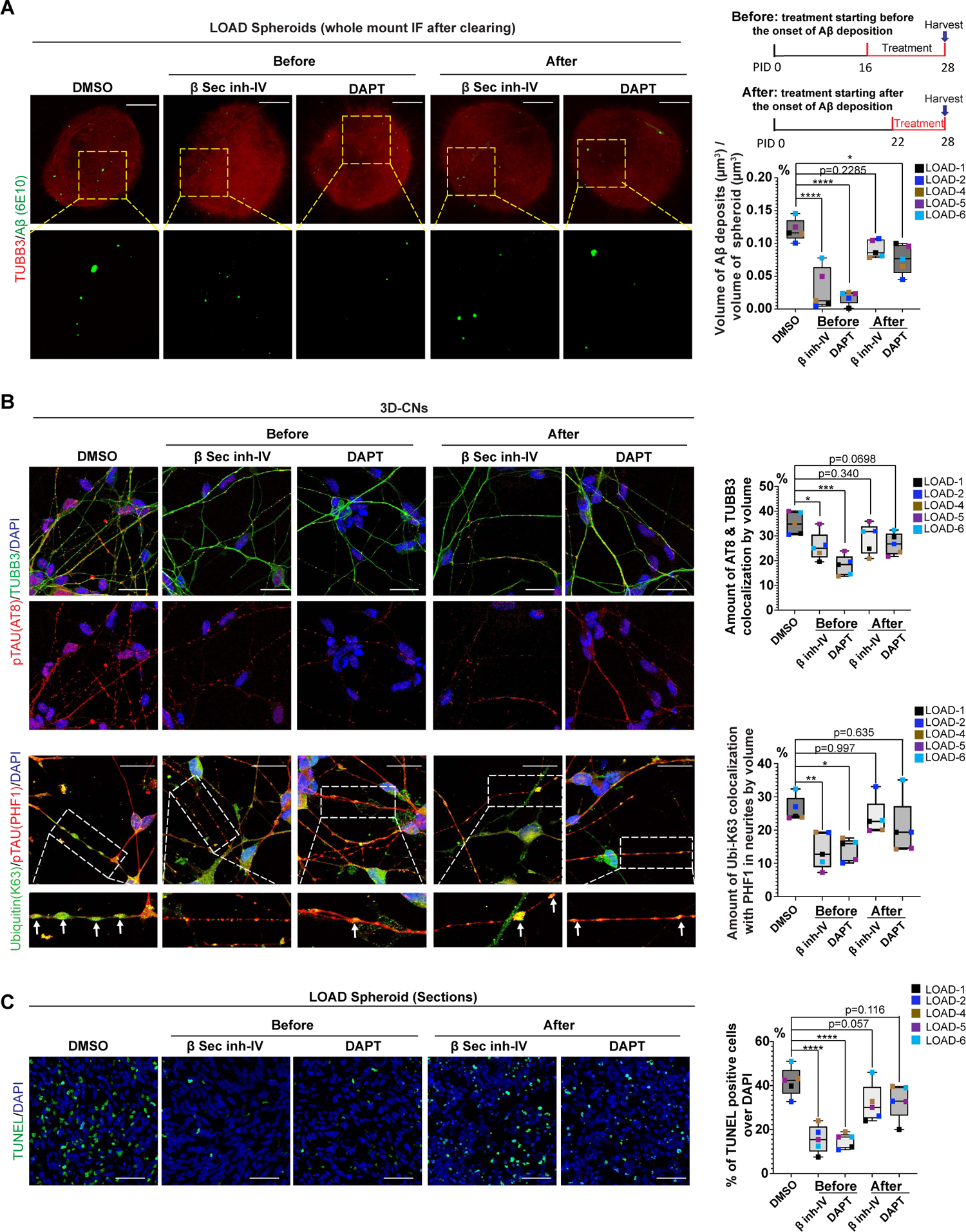
Effects of inhibiting APP processing in LOAD 3D-CNs and spheroids on tauopathy and neurodegeneration. **A.** LOAD spheroids were treated with β-secretase inhibitor IV or DAPT starting at PID16 (before the observed onset of Aβ deposition) or PID22 (after the onset of Aβ deposition) and Aβ deposition was examined at PID28 by whole mount immunostaining using Aβ antibody (6E10). Scale bar: 500 µm. **B.** Tau phosphorylation (AT8, top panel) and K63-ubiquitin/p-tau (PHF1) colocalization (bottom panel) were examined in PID28 LOAD 3D-CNs treated with β-secretase inhibitor IV or DAPT starting at PID16 or PID22. Arrows highlight swelled tau blebs in the bottom panel. Scale bar = 25 µm. **C.** TUNEL staining of the sections of PID28 LOAD spheroids that were treated with β-secretase inhibitor IV or DAPT starting at PID16 or PID22. Scale bar: 50 µm. For quantifications in A, B, and C, n = 5 independent LOAD patients. Imaris was used to calculate the volumes (µm^3^) or numbers of objects marked by different antibodies. Adjusted p-values were calculated by two-way ANOVA with Šídák’s multiple comparisons test. *adjusted p < 0.05; **adjusted p < 0.01; ****adjusted p < 0.0001.

### LOAD spheroids exhibit distinct transcriptomes and inhibition of retrotransposable element (RTE) dysregulation suppresses LOAD pathologies

To deduce transcriptional patterns associated with LOAD, we conducted RNA sequencing (RNA-seq) on PID25 spheroids reprogrammed from five independent LOAD fibroblast lines and five sex- and age-matched HC fibroblast lines. Principle component analysis indicated a clear separation of samples based on disease status (HC vs LOAD) (Fig. S7A). By comparing protein-coding gene expression in LOAD and HC spheroids, we identified 846 differentially expressed genes (DEGs) (P < 0.05, |log_2_FC| > 0.58) (Fig. 6A). Notably, several MMP (M10 matrix metallopeptidases) genes, including *MMP1*, *MMP3* and *MMP8* were among the most upregulated genes. MMPs are known to be linked to extracellular protein degradation including Aβ^49^, while MMPs upregulated in AD may also confer neuroinflammation, synaptic dysfunction, and neuronal death^50, 51^. Notably, gene ontology analysis revealed that upregulated DEGs in LOAD spheroids were significantly enriched for pathways associated with immune response and inflammatory responses, while downregulated DEGs were associated with nucleosome assembly, telomere organization, and memory (Fig. 6B).

**Fig. 6.**
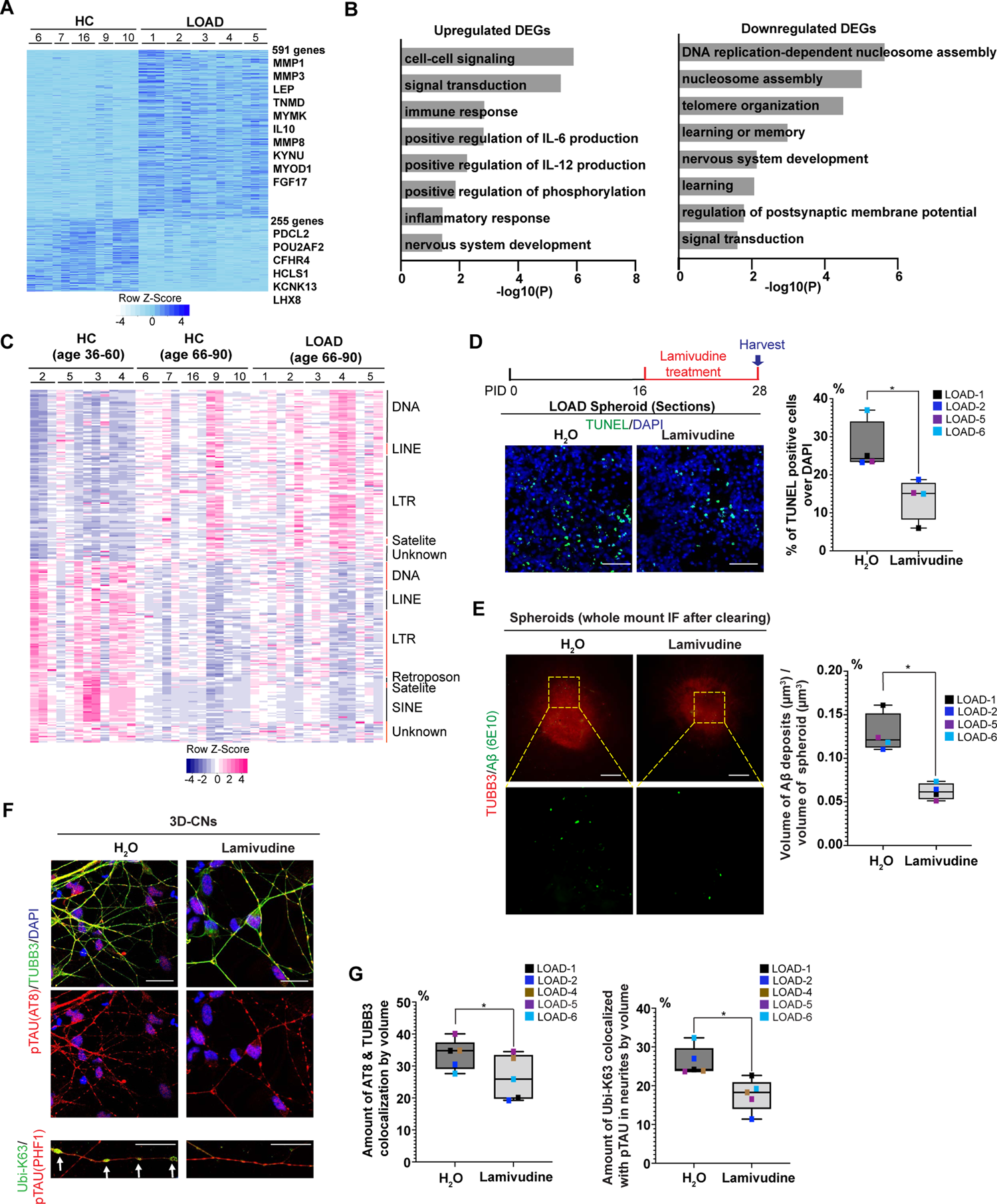
Differential gene expression in LOAD spheroids and Lamivudine, a reverse transcriptase inhibitor, ameliorates LOAD neuropathologies. **A.** Heatmap of gene expression levels of significant (p < 0.05, |log2FC| > 0.58) differentially expressed genes (DEGs) in PID25 LOAD and HC spheroids by RNA-seq. A total of 28 samples from 10 independent individuals (5 for LOAD and 5 for HC) were used for analysis. **B.** Top Gene Ontology (GO) terms associated with upregulated (left) and downregulated (right) DEGs analyzed by DAVID analysis. **C.** A heatmap of significant (p < 0.05, |log2FC| > 0.58) differentially expressed transposable elements (DETEs) in PID25 LOAD and HC spheroids analyzed by RNA-seq. **D.** Top: Schematic of lamivudine treatment for inhibiting RTE formation in LOAD spheroids. Bottom images: TUNEL staining of the dead cells (green color) in lamivudine- or H_2_O-(control) treated spheroids (PID28). Scale bar: 50 µm. **E.** Representative whole mount immunostaining images of Aβ deposition in lamivudine and H_2_O-treated spheroids at PID28. Scale bar: 500 µm. **F.** p-Tau labeled by AT8 antibody (top panel) and co-localization of p-tau (PHF1) and K63-ubiquitin (bottom panel) in lamivudine or H_2_O-treated LOAD 3D-CNs at PID28. Arrows highlight the dystrophic swelled tau blebs. Scale bar = 25 µm. **G.** Quantification of F. Percentages of p-tau (AT-8) colocalized with TUBB3 (left) and K63-linked ubiquitin colocalized with p-tau (PHF1) in neurites (right) of LOAD 3D-CNs treated with lamivudine or H_2_O. For quantifications: n = 4-5 LOAD patient lines in D, E, and G. *p < 0.05 was calculated by paired t-test.

While inflammation can result from various factors, it has been shown that activation of retrotransposon elements (RTEs) in aging and late-onset diseases may trigger cell-intrinsic inflammation^52, 53^. Suppression of RTE ameliorates age-associated inflammation^52, 54^ and reduces tau activiation and tau-induced neurotoxicity in tau transgenic *Drosophila*^55^. Consequently, we examined whether reprogrammed spheroids exhibited age-dependent changes in the expression of transposon elements (TEs). TE analysis from RNA-seq datasets of HC spheroids divided by age (36-60 years vs. 66-90 years) revealed differentially expressed TEs (DETEs) between age groups, with TEs linked to the older age group also enriched in LOAD spheroids (Fig. 6C). To determine whether inhibiting RTE expression could influence AD-associated phenotypes, we treated LOAD spheroids and 3D-CNs with the reverse transcriptase inhibitor lamivudine (3TC). Remarkably, 3TC treatment from PID16 to PID28 significantly reduced neuronal cell death in LOAD spheroids compared to the control H_2_O treatment (Fig. 6D). Additionally, 3TC considerably decreased Aβ formation and tau dysregulation, as evidenced by reduced Aβ deposition, p-tau, and K63-linked ubiquitin-positive tau signals (Fig. 6E, F, and G). This protective effect was not due to the loss of neuronal identity with 3TC treatment, as assessed by qPCR (Fig. S7B). These results demonstrate that perturbing RTE synthesis conferred protective effects in LOAD neurons and highlight the overall potential of using the patient-specific LOAD neurons to track changes in AD phenotypes for chemical interventions.

## Discussion

Our findings highlight the significant advance in modelling AD through 3D neuronal reprogramming: 1) the age-maintained patient-derived neurons recapitulate hallmark neuropathological features of AD, including extracellular Aβ deposition, tauopathy, and neurodegeneration, 2) the capacity to endogenously capture neuronal aspects of AD pathology without requiring additional cellular insults or transgenes, and 3) the potential to model late-onset neuropathology of LOAD. The finding that 3D-reprogrammed neurons derived from LOAD patients, without genetic mutations linked to ADAD, exhibit neurodegeneration-related phenotypes underscores the efficacy of 3D neuronal reprogramming in capturing AD-associated traits resulting from the donor’s genetic background and epigenetic signatures. Although the current study provides evidence that inhibiting APP processing or dampening RTE promoted the survival of reprogrammed LOAD neurons, identifying other factors contributing to the late-onset degeneration of LOAD neurons would help inform strategies to slow neurodegeneration. Consequently, future research using whole genome-sequencing and chromatin profiling would facilitate the identification of genetic and epigenetic changes underlying the neurodegeneration phenotype observed in patient-specific neurons. The 3D neuronal reprogramming could serve as a platform to investigate genetic risk factors that may function in neurons, contributing to individual variability and vulnerability to neuropathology of AD. It is also noteworthy that Aβ deposits and p-tau signals correlate with aging even in neurons derived from healthy individuals (Fig. S4A and Fig. S5A). This observation emphasizes the potential of using 3D neuronal reprogramming to explore how neuronal aging generally increases susceptibility to late-onset neurodegeneration.

Here, we highlight the significant differences of 3D direct reprogramming modeling approach provided in this study compared with animal and human cellular models based on ADAD-associated mutations or 2D direct reprogramming approaches. While mouse models based on overexpressing gene mutations associated with ADAD offer valuable insights into Aβ plaque formation^56^, they show limited tau pathology. It typically requires additional expression of human tau harboring mutations associated with primary tauopathy^57–59^ or seeding mutant human *APP* knock-in mouse brain with human AD brain-derived tau to develop the tau pathology^60^. As for human neurons, previous studies focused on overexpressing APP and Presenilin containing ADAD mutations in human stem cell lines^3^, in which they provided evidence that these cell lines developed Aβ accumulation and tauopathy^3^.

Generation of age-equivalent, AD patient-derived neurons in 2D-culture environment has been recently carried out by using Ngn2/Ascl1-based direct conversion approach, which revealed AD patient-derived neurons exhibited age-dependent instability of mature neuronal fate^13^, increased post-mitotic senescence and pro-inflammatory signature^61^. Here, we focused on the feasibility of endogenously capturing critical neuropathological features of AD in patient-derived neurons by developing 3D-direct reprogramming approaches. In this 3D culture system, AD patient-derived neurons not only exhibited robust extracellular Aβ deposition and tau dysregulation, but also developed severe neurodegeneration. Moreover, we validated that inhibiting APP processing in patient-derived neurons and spheroids alleviates tau pathology, and importantly, it also reduces the neurodegeneration phenotype. However, this effect was only minimal when APP processing was inhibited after LOAD neurons had already started forming Aβ deposits. Therefore, AD modelling through 3D neuronal conversion can serve as a platform to study the interplay between Aβ deposition, tau dysregulation and neurodegeneration, and identify gene targets or small molecules, as exemplified by 3TC experiments, that can perturb the disease progression.

Results in this study also demonstrate the sufficiency of directly reprogrammed patient neurons to capture key neuronal features of AD which prompts new avenues for studying how these phenotypes may change when AD neurons interact with other cell types in the context of aging and disease. In this regard, aging may impact glial cells such as astrocytes^62, 63^ and microglia^64^. Thus, robust direct reprogramming methods for generating aged astrocytes and microglia would enable the study of neuron-glia interactions during the aging process and age-associated disease progression. In conclusion, our study demonstrates the effectiveness and sufficiency of 3D-cultured, patient-derived neurons through miRNAs-mediated direct reprograming for modeling critical neuropathological hallmarks of AD. This approach presents opportunities to investigate molecular events intrinsic to neurons that drive AD-associated neurodegeneration and may serve as a patient-specific neuron platform for testing various compounds or target genes for personalized therapeutic interventions.

## Methods

### Fibroblast cell lines

Adult dermal fibroblast cell lines from familial AD patients were obtained from University College London (UCL33, UCL53 and UCL803) and Coriell Institute for Medical Research (AG06840). Adult dermal fibroblast cell lines from biomarker positive late-onset AD patients were acquired from Washington University Knight Alzheimer’s Disease Research Center (ADRC), which includes FA15-552, FA12-453, FA17-623 and FA18-633. Two additional skin fibroblast cell lines from late-onset Alzheimer’s disease patients were purchased from Coriell (AG05810 and AG06869). The skin fibroblast cell lines derived from healthy control individuals were obtained from Washington University Knight ADRC (biomarker negative: FA18-634 and FA12-463) and Coriell (AG08260, AG04453, AG04060, AG05838, AG13246, AG11020, AG13369, AG08379, AG14251, AG04356, GM02171, AG07307 and AG11246). Detailed information of the list of fibroblast cell lines used in this study is provided in Table S1. All fibroblast cell lines were cultured in fibroblast medium (FM) containing 15% fetal bovine serum (FBS): Dulbecco’s Modified Eagle Medium (DMEM, Invitrogen) with high glucose containing 15% FBS, 0.1% 55 mM beta-mercaptoethanol, 1% 1M HEPES buffer, 1% nonessential amino acids, 1% sodium pyruvate, 1% GlutaMAX and 1% penicillin/streptomycin solution (all from Gibco). Cell culture medium was routinely checked to ensure there was no mycoplasma contamination. Cells were maintained up to 15 passages.

### Lentiviral production

Lentiviruses were generated as previously described^26^. Briefly, the second-generation lentiviral system for production was used by transfecting the transfer plasmid, packaging plasmid (psPAX2) and envelope plasmid (pMd2G) in HEK293LE cells. 60-70 hours after transfection, the supernatant containing lentivirus was filtered through a 0.45 µm PES membrane and concentrated at 70,000 g for 2h at 4°C. Viral pellets then were resuspended in PBS, aliquoted and stored at −80°C until used for transduction (viruses can be stored in −80°C for up to a year).

### 3D direct neuronal reprogramming

The 3D direct neuronal reprogramming protocols were established based on modifications of our previously published 2D direct reprogramming protocol^23, 26^. Briefly, human fibroblasts were spin-infected (1000g for 30 mins at 37°C) with lentiviral cocktail containing rtTA, pTight-9-124-BclxL, MYT1L and NEUROD2. After 16-18 hours incubation at 37°C, the infected cells were washed once with PBS and fed with fresh fibroblast medium containing 10% FBS and 1 μg/ml doxycycline (Sigma, cat. #D9891). The fibroblast medium was changed every other day with 10% FBS, 1 μg/ml doxycycline and antibiotics. On day 7, cells were trypsinized, resuspended in transiting medium and counted for replating.

For thin gel culture, 0.1×10^6^ cells resuspended in 100 μL transiting medium containing 15% Matrigel (Corning, cat. # 354230) were immediately transferred to each well of an optically clear 96-well plate (Perkin Elmer, cat. # 6055302) to form a thin gel. The plates were incubated at 37°C overnight to let the gel solidify and form a thin layer (∼100 μm) matrigel/cell mixture at the bottom of the well. The next day (day 8), 200 μL pre-warmed Neurobasal-A medium was added to the thin gel.

For spheroid culture, 0.5×10^6^ cells were centrifuged at 400g for 2.5 min at room temperature to form a cell pellet. The cell pellet was transferred to the center of a transwell insert (Corning, cat. # 353095) in a 24 well plate by using a wide-bore tip (200μL, USA Scientific. cat. # 1011-8410) to form a spheroid. 250 μL transiting medium was added to the bottom of the well and the plate containing spheroids was placed in 37°C incubator overnight. On day 8, the spent transiting medium in the bottom well of transwell plates was replaced by 250 μL fresh Neurobasal-A medium. 100 μL of 50% Matrigel diluted in Neurobasal-A medium was added into the transwell insert. After incubation at 37°C for 3 hours to overnight to let the Matrigel solidify and form a “Matrigel blanket”, 250 μL fresh Neurobasal-A medium was added on top of the “Matrigel blanket”.

After replating, half media changes (Neurobasal-A medium) were performed every other day for both thin gel culture and spheroid culture. On day 14 and after, Neurobasal-A medium was replaced by Brainphys neuronal medium for half media changes. Puromycin (3 µg/mL) was added to the medium from PID3 to PID14 and G418 (geneticin, 300 µg/ml) was added to the media at PID5-PID10 for selection. 1 μg/ml doxycycline was added until PID30.

Transiting medium: Neurobasal A medium mixed with fibroblast medium (10% FBS) at ratio of 1:1. No antibiotics were added to transiting medium.

Neurobasal-A medium: Neurobasal-A medium (Gibco, cat. # 10888022) containing B-27 plus supplement (Gibco, cat. # A3582801), GlutaMAX (Gibco, cat. # 35050061) and supplemented with doxycycline (1 μg/mL), valproic acid (1 mM), dibutyl cAMP (200 μM), BDNF (10 ng/ mL), NT-3 (10 ng/ mL), retinoic acid (1 μM), ascorbic acid (200 nM), and RVC (RevitaCell Supplement, 1×).

BrainPhys neuronal medium: BrainPhys neuronal medium (Stemcell Technologies, cat. # 05790) containing NeuroCult SM1neuronal supplement (Stemcell Technologies, cat. # 05711), N2-A supplement (Stemcell Technologies, cat. # 07152) and following supplements: doxycycline (1 μg/mL), valproic acid (1 mM), dibutyl cAMP (200 μM), BDNF (10 ng/ mL), NT-3 (10 ng/ mL), Retinoic acid (1 μM), and ascorbic acid (200 nM).

### β-secretase inhibitor IV, DAPT and lamivudine (3TC) treatment

β-secretase inhibitor IV (cat. # 565788), DAPT (cat. # D5942), and lamivudine (3TC) (cat. # PHR1365) were purchased from Millipore Sigma. To make the stock solution, 5 mM β-secretase inhibitor IV and 40 mM DAPT were dissolved in DMSO and 100 mM 3TC was dissolved in sterile water. For thin gel culture, 2.5 μM β-secretase inhibitor IV, 20 μM DAPT, or 100 μM 3TC was added to the fresh media and used for half media changes starting at PID16 or PID22. For spheroid culture, 5 μM β-secretase inhibitor IV, 40 μM DAPT, or 100 μM 3TC was added to the fresh media and used for half media changes starting at PID16 or PID22.

### Fixation of 3D cultures, OCT embedding and sectioning of neuronal spheroids

Fixation: For thin gel culture, 100 μL 4% (wt/vol) paraformaldehyde (PFA) solution was added to each well; For spheroid culture, 250 μL 4% (wt/vol) PFA solution was added in the bottom well and another 250 μL on the insert. After incubating overnight at room temperature, the PFA solution was removed and the 3D cultures were washed with PBS once. Plates containing fixed 3D cultures were filled with PBS and sealed with parafilm to prevent evaporation. The fixed 3D cultures can be stored at 4°C for 6-12 months.

OCT embedding and sectioning of neuronal spheroids: Fixed neuronal spheroids were soaked in 30% sucrose (w/v) overnight at 4°C. Neuronal spheroids were manually detached from the transwell insert, embedded in OCT compound (Tissue Tek), and followed by snap-frozen in liquid nitrogen to form OCT blocks. The OCT embedded spheroids were cryo-sectioned at 20 µm and mounted on positively charged Superfrost slides (VWR). Slides can be stored at −80°C until further use.

### Immunofluorescence staining, 3D reconstruction, and quantification

Immunofluorescence staining of thin gel: The fixed thin gel cultures were permeabilized in PBST (0.2% (vol/vol) TritonX-100 in PBS) at room temperature for 1h and blocked with blocking buffer (5% (wt/vol) BSA (Sigma, cat. # A2153), 2% (wt/vol) normal goat serum (Sigma, cat. # G9023), and 0.2% (vol/vol) TritonX-100) overnight at 4°C. The blocking buffer was removed from thin gel culture plates and primary antibodies diluted in blocking buffer were added to the plate followed by incubation overnight at 4°C with gentle rocking. After washing 5 times with PBST, the cells were then incubated with secondary antibodies diluted in blocking buffer at room temperature for 3-5 hours or 4°C overnight. Cells were washed 5 times with PBST, incubated with DAPI (Sigma) for 20 min at room temperature if needed and sealed with a drop of anti-fade prolong gold reagent (Life technology). Plates were sealed with parafilm and stored at 4°C before imaging. Immunofluorescence images were taken on a Leica SP5X white light laser confocal system.

Immunofluorescence staining of sections from spheroids: Sections were permeabilized in PBST for 10 min followed by incubating in blocking buffer for 30 min at room temperature. The sections were incubated with primary antibodies in blocking buffer at 4°C overnight, then washed three times with PBST. The sections were incubated with secondary antibodies in blocking buffer for 30-60 min at room temperature, followed by three washes with PBST. The sections were then stained with DAPI for 10 min at room temperature and mounted in anti-fade prolong gold reagent (Life technologies).

Immunofluorescence images were taken on a Leica SP5X white light laser confocal system. Whole mount immunofluorescence staining of spheroids: For whole-mount immunostaining of neuronal spheroids, 3D cell culture clearing kit (Abcam, cat# ab243299) was used with modifications. Briefly, fixed spheroids on transwell inserts were permeabilized by sequential dehydration and rehydration using different concentrations of ethanol. Specifically, spheroids were sequentially soaked in 50% ethanol in PBS, 80% ethanol in PBS, 100% ethanol, 80% ethanol in DMSO, 80% ethanol in PBS, 50% ethanol in PBS, and PBS (5 min each, room temperature). Spheroids were incubated in the Tissue Clearing Penetration buffer (from the kit) for 30-60 mins at room temperature followed by blocking overnight in blocking buffer. Then the spheroids were incubated with primary antibodies diluted in blocking buffer at 4°C for 24-48 hours with gentle rocking, followed by washing 5 times with PBST, 10 mins each. Fluorescent secondary antibodies diluted in blocking buffer were added to the spheroids and incubated overnight at 4°C. After 5 washes with PBST, the 3D Cell Culture Clearing Reagent (from the kit) was added to completely cover the spheroids and incubated for at least 1 hour at room temperature with gentle rocking. The spheroids should look transparent under light microscope. The spheroids mounted in this clearing reagent can be stored in 4°C for at least 6 months with proper parafilm sealing of the plates and avoiding light. Immunofluorescence images were taken on an upright Nikon A1RHD25 MP multi-photon microscope.

For the list all the antibodies and dilution ratios used in this study, please see Table S2. For quantification of the total volume of Aβ deposits, volume and area of the spheroids, co-localization analysis in tau related pathologies, and number of TUNEL positive cells or DAPI positive cells, the “surface” module in Imaris was used for 3D reconstruction and quantification. For measuring the intensity of neurite outgrowth in spheroids, Image J was used by measuring the mean intensity at each region (distal, middle, and proximal).

### FRET live imaging

Tau RD P301S Ruby2 and tau RD P301S clover plasmids were constructed as previously described^39^. Lentiviral FM5-Ruby2 or FM5-Clover plasmid drives the tau repeat domain (RD, aa246 to 378) with the disease-associated P301S mutation under the human ubiquitin C promoter. The lentiviruses to express tau RD (P301S)-Ruby2 and RD (P301S)-clover were transduced into reprogrammed cortical neurons at PID19 using Polybrene (Sigma-Aldrich, cat. # H9268). The next day, the media containing lentiviruses were removed and fresh Brainphys neuronal medium was added to the cells. Live cell imaging was performed at PID25 or PID28 using Leica SP5X white light laser confocal system. To detect FRET, cells were illuminated with 488 nm light and emission was captured between 620 to 700 nm. To avoid false positive signal in the FRET channel, cells in separate wells were transduced with single reporters (RD (P301S)-Ruby2 or RD (P301S)-clover only) to set imaging thresholds.

### PCR analysis of 3-repeat (3R) and 4-repeat (4R) tau mRNA levels

Semi-quantitative PCR for analyzing 3R and 4R mRNA expression was carried out as previously described^16^. Briefly, cDNAs that were reverse transcribed from RNAs of LOAD and HC 3D-CNs were used for PCR amplification using primers ^3^ (forward 5’-AAGTCGCCGTCTTCCGCCAAG-3’; reverse 5’-GTCCAGGGACCCAATCTTCGA-3’) flanking exon 10 of tau. The PCR products were run on a 2% agarose gel and images were taken on a Chemidoc MP imaging system (BIO-RAD). Image J was used to analyze the pixel intensity of 3R (288 bp) and 4R (381 bp) bands.

### Electrochemiluminescence assay for Aβ42 detection

For Aβ42 detection, V-PLEX Plus Aβ Peptide Panel 1 (6E10) Kit (Meso Scale Discovery (MSD), cat. # K15200E0) was used as previously described ^65^. 2-3 neuronal spheroids from the same cell line were pooled together and lysed in RIPA buffer using a sonicator. Spheroids reprogrammed from 4 ADAD fibroblast lines and 4 age- and sex-matched healthy control lines were used for the experiment. Aβ42 levels were measured using the MESO QuickPlex SQ 120 (multiplexing imager, MSD) following the manufacturer’s instructions. Aβ42 level was normalized to the total protein level in each sample.

### RNA preparations and quantitative PCR (qPCR)

Total RNA was extracted from thin gel cultures or neuronal spheroids using TRIzol (Invitrogen) followed by RNeasy Micro Kit (Qiagen, cat. # 74004). Briefly, cells in thin gel or spheroids were lysed by TRIzol at room temperature for 5 mins. 0.2 mL of chloroform was added per 1 mL of TRIzol reagent and incubated for 2-3 mins at room temperature. The TRIzol/chloroform mixtures were centrifuged for 5 mins at 12,000g at 4°C. Aqueous phase was transferred to a fresh microcentrifuge tube and mixed with 1 volume of 70% ethanol. Up to 700 μL of clear lysates/70% ethanol mixture was applied to RNeasy MinElute column. Washing and elution were performed following the instructions from the RNeasy Micro Kit.

Reverse transcription was performed using SuperScript IV First Strand Synthesis SuperMix (Invitrogen, cat. # 18090050) according to the manufacturer’s protocol. qPCR was performed with fast SYBR Green PCR Master Mix (Applied Biosystems, cat. # 4385612) using the StepOne Plus Real-Time PCR System (AB Applied Biosystems, Germany). All samples were run with two technical replicates. All absolute data were normalized to GAPDH and fold change was calculated based on the 2^-ΔΔCT^ method. The sequences of primers used for qPCR are as listed in Table S3.

### Electrophysiology

Whole-cell patch-clamp recordings were performed as previously reported ^29, 66, 67^. Briefly, reprogrammed cortical neurons were transduced with pSynapsin-RFP at PID3 to label neurons. At PID 14, human astrocytes (Sciencell, cat. # 1800) were added on top of the neurons for co-culturing.

Brainphys neuronal medium and Astrocyte medium (ScienCell, cat. # 1801) were mixed at 1:1 for half media changes at PID14 and after. Whole cell patch clamp was performed at PID30. During recording, reprogrammed cortical neurons were transferred to a recording chamber with continuous perfusion (2 ml/min, 32°C) of oxygenated, regular aCSF (in mM: 125 NaCl, 25 glucose, 25 NaHCO_3_, 2.5 KCl, 1.25 NaH_2_PO4, 2.5 CaCl_2_, 1.2 MgCl_2_ equilibrated with 95% oxygen, 5% CO_2_ plus 2.5 CaCl_2_, 1.2 MgCl_2_; 310 mOsm). Individual neurons were visualized and identified by IR-DIC microscopy (Nikon FN1 microscope and Photometrics Prime camera). Borosilicate glass pipettes (World Precision Instruments, Inc) with open tip resistance of 3-7 MΩ were used for whole-cell recording. Pipettes were filled with potassium gluconate solution containing the following (in mM: 120 K-Gluconate, 10 KCl, 2 EGTA, 10 HEPES, 2 MgATP, and 0.3 Na_2_GTP; pH 7.25 with KOH; 280-290 mOsm). Recordings were acquired using pCLAMP 10.4 software with a MultiClamp 700B amplifier and Digidata 1550 digitizer (Molecular Devices). Action potentials were elicited by current injection with 8 pA increments in current-clamp mode. From voltage-clamp mode, voltages step (100 ms) in increments of 10 mV were applied from a holding potential of −70 mV to monitor currents. Data were acquired at 5 kHz sampling rate and filtered at 2 kHz.

### SYTOX assay and TUNEL assay

SYTOX: for thin gel cultured cortical neurons, 0.1 μM SYTOX green nucleic acid stain (Invitrogen, cat. # S7020) and 1 μg/mL of Hoechst 33342 (ThermoFisher Scientific, cat. # H3570) were added into cell media. Samples were incubated for at least 30 mins at 37°C prior to the live cell imaging. Images were captured with a GE InCell 200 fluorescence microscope and analyzed by InCell investigator and developer image analyses software. Quantifications were performed by counting the percentage of SYTOX positive cells over Hoechst.

TUNEL assay: DeadEND Fluorometric TUNEL system kit was purchased from Promega (cat. # G3250) and TUNEL assay was performed according to the manufacturer’s description. Briefly, the cryosections of neuronal spheroids were permeabilized by 0.2% TritonX-100 in PBS for 5 mins followed by two 5 mins washes in PBS. The sections then were equilibrated in equilibration buffer for 5-10 mins at room temperature and labeled by TdT reaction mix for 1 hour at 37°C in a humidified chamber. The sections were immersed in 2x SSC for 15 min to stop the reaction, followed by three 5 mins washes in PBS. The sections then were counterstained with DAPI and mounted in anti-fade prolong gold reagent (Life technologies). Immunofluorescence images were taken on a Leica SP5X white light laser confocal system.

### Electron microscopy

High-pressure freezing: Reprogrammed cortical neurons were seeded and grown on 3 mm sapphire discs (Leica) until PID30. The 100 µm cavity of A-type specimen carrier (Leica) was filled with 20% BSA in culture medium and sapphire discs with cells facing the cryoprotectant were transferred on top of the carrier. A 200 µm thick ring for alternative spacing was placed on top of the assembly that was then vitrified using high-pressure freezing machine (Leica EM ICE).

Neuronal spheroids (PID28) were fixed for one hour in 4% PFA in PBS, rinsed three times in PBS and then embedded in 4% agarose in PBS. Embedded specimens were cut into 200 µm sections using a vibratome (Leica VT1200S). Sections were placed in a 300 µm cavity of a B-type specimen carrier (Leica) filled with 20% BSA in PBS. The assembly was covered with a flat side of another B-type carrier and specimens were vitrified using a high-pressure freezing machine (Leica EM ICE).

Freeze-substitution: The frozen specimens were stored in liquid nitrogen until further processing. The freeze-substitution process was performed using a cocktail of 0.1% uranyl acetate in acetone and EM AFS2 machine with FSP robot for automated reagent handling (Leica). Briefly, samples were kept for 50 hours at −85°C and then warmed up over a period of 11 hours to −50 °C. At −50 °C samples were washed four times in ethanol for 30 minutes each before gradual infiltration with HM20 resin (Electron Microscopy Sciences). Polymerization of the resin was done with a UV light source and was carried out at −50 °C for 48 hours followed by post-polymerization at room temperature for 2 days.

Immunogold labelling: Thin sections of 70-80 nm were cut with an ultramicrotome (Leica, UC7) and placed on nickel grids (Ted Pella). Blocking was performed with 1% BSA in PBS and sections were incubated with primary antibody (Aβ antibody, 6E10) prepared at 1:20 dilution in the blocking buffer overnight at 4 °C. Grids were washed five times for five minutes each in the blocking buffer. Following this, sections were incubated with 12 nm gold-conjugated donkey anti-mouse (Jackson ImmunoResearch Labs, cat. # 715-205-150) secondary antibody (1:30 dilution in the blocking buffer) for 1 hour at room temperature. Sections were then washed in PBS and ultrapure water, dried and imaged on a TEM (Jeol JEM-1400 Plus) at 120 kV.TEM images were acquired using an AMT Nanosprint15-MkII sCMOS camera.

### RNA-seq cell collection, sequencing, data processing, and analysis

For comparing the transcriptome between LOAD vs HC spheroids, 3 biological replicates from 5 independent cell lines per disease group were used for RNA sequencing. Each replicate consisted of RNAs pooled from 2-3 spheroids. For RNA-seq of thin gel samples harvested at different time points, 2 biological replicates from one control cell line and one LOAD cell line were used for RNA-seq. Each biological replicate contained RNAs pooled from 5-10 wells of a 96 well plate. RNA samples that passed quality control were submitted to Genome Access Technology Center at Washington University for library preparation and sequencing. NovaSeq6000 using SMARTer 150PE with 30 million reads per sample input was used for sequencing. FastQC^68^ was used to determine sequencing quality and identify adapter contamination. Raw fastq files were trimmed using Cutadapt v3.2^69^ for adapter contamination and low sequencing quality. Trimmed sequences were mapped to GRCh38 genome reference using STAR v2.7.10a^70^. Raw gene counts were extracted using STAR option --quantMode GeneCounts. Differential gene expression analysis was performed using RUVSeq^71^ and DESeq2^72^ normalizing to sequencing depth and removing unwanted variation.

### Transposable Elements (TE) analysis

Trimmed reads from RNA-seq data analysis aligned to GRCh38 human genome using the STAR v2.7.10a with the additional parameters as recommended in the TE transcripts manual: -- winAnchorMultimapNmax 100 and –outFilterMultimapNmax 100. TE transcript counts were generated based on RepeatMasker database using TEtranscripts v2.2.3^73^. Differentially expressed TEs at the subfamily level between two different conditions were identified by DESeq2.

### Mass spectrometry analyses of Aβ

Aβ38, 40, 42 were quantified by mass spectrometry as previously described with some modifications^74, 75^. 0.5-1mL of spheroid media was immunoprecipitated with monoclonal anti-Aβ mid-domain antibody (HJ5.1, anti-Aβ_13–28_) conjugated to Sepharose beads (30 μL, 3 mg/mL). Aβ was digested on beads with 50 μL of LysN (0.25 ng/μL) in 25 mM triethyl ammonium bicarbonate (TEABC). Digests were desalted by C18 TopTip (Glygen). Before eluting samples, 3% hydrogen peroxide and 3% formic acid (FA) in water was added onto the beads, followed by overnight incubation at 4 °C to oxidize the peptides containing methionine. The eluent was lyophilized and resuspended in 25 μL of 2-10% FA and 2-10% acetonitrile and 20 nM BSA digest prior to mass spectrometry analysis on nanoAcquity UPLC system (Waters) coupled to Orbitrap Fusion mass spectrometer (Thermo Scientific) operating in selected reaction mode. Concentrations of each peptide were estimated using internal standards containing Aβ38, 40, 42 uniformly labeled with ^15^N (0.01875, 0.125 and 0.0125 ng/μL, respectively. rPeptide).

### Statistics

For quantifying immunostaining data from thin gel culture, 3 to 4 confocal images were taken from different areas of each well for 2 wells of cell culture per each cell line, with 3 to 6 independent cell lines per disease group (AD vs HC) being used for analysis in each experiment. For quantifying whole mount immunostaining data in spheroid culture, 1 confocal image was taken from the whole area of a spheroid for 2-3 spheroids per cell line, and 3 to 6 independent cell lines per disease group were used for analysis. For quantifying TUNEL data using spheroid sections, 3 to 4 confocal images were taken from different area on each section for 3 sections covering different planes of the spheroid, with 2-3 spheroids of cells per cell line, and 3 to 6 independent cell lines per disease group were used for analysis. For harvesting RNA and qPCR analysis, 2-3 spheroids per cell line were pooled together for harvesting one RNA sample, with 2-3 RNA samples per cell line, and at least 3 independent cell lines per disease group or per treatment were used for the experiments. An extra technical duplicate for each sample was used for qPCR analysis. Statistical analyses were performed in GraphPad Prism (version 9.5.1(528)). Specific “n” information and statistical analysis method for each assay can be found in the figure legend. Generally, two-tailed paired or unpaired Student’s t-tests were performed for datasets containing two groups. One-way or two-way ANOVA analyses were used for datasets containing more than two groups.

## Acknowledgements

We acknowledge the assistance of Washington University Center for Cellular Imaging (WUCCI) in electron microscopy studies, the Genome Technology Access Center (GTAC) at Washington University for RNA-sequencing experiments, Rama Krishna Koppisetti and Chloe He for helping the Mass spectrometry analyses of Aβ, Drs. Jason Ulrich and Chanung Wang for sharing the brain sections from 5XFAD mouse and other reagents, Dr. Virginia Man-Yee Lee at University of Pennsylvania for sharing the GT-38 tau antibody, Joshua D. Beaver from UT Southwestern Medical Center for technical support of FRET assay, Dr. John M Sedivy from Brown University for suggestions of TE analysis, Dr. Irving Boime for editing manuscript, Luorongxin (Mini) Yuan and Kyle Burbach for drawing the cartoon of spheroid and thin gel culture. We recognize BioRender.com for generating some cartoons in Fig.1. This study was supported by the following programs, grants and fellowships: Farrell Family Fund for Alzheimer’s Disease (A.S.Y., C.M.K., and D.M.H.); Cure Alzheimer’s Fund (A.S.Y.); Centene Fund (A.S.Y., C.M.K., and D.M.H.), NIH/NIA RF1AG056296 (A.S.Y.); NIH/NINDS R01NS107488 (A.S.Y.); NIH/NIA R01AG078964 (A.S.Y., C.M.K.); NIH NIA AG066444 (C.M.K.); Mallinckrodt Scholar Award (A.S.Y.); Cure Alzheimer’s Fund (R.E.T.); The Children’s Discovery Institute of Washington University and St. Louis Children’s Hospital CDI-CORE-2015-505 and CDI-CORE-2019-813 (WUCCI); Foundation for Barnes-Jewish Hospital 3770 and 4642 (WUCCI); Knight Alzheimer Disease Research Center (AG066444).

## Author contributions

Z.S. and A.S.Y. designed experiments and wrote the manuscript. Z.S. performed experiments and analyzed data. J.S.K. analyzed the RNA-seq data in figure 6. Y.R. performed TUNEL experiments for 3TC treated spheroids. S.C. performed some of the 3D-reprogramming experiments. K.C. performed LGE analysis and generated heatmaps in Figure S1. X.L. and S.J.M. performed the electrophysiology and analyzed the data. C.K.W. edited the manuscript. H.K. and J.K. performed electrochemiluminescence assay for Aβ42 detection and analyzed data. S.S. and J.A.F. performed the electron microscopy assay. C.V. and M.I.D. provided the constructs and guidance for tau FRET reporter assays. H.H. provided some of ADAD patient fibroblast lines. C.M.K provided some of the LOAD patient fibroblast lines used in the study. R.J.B. and C.S. performed mass spec for measurement of Aβ species. R.E.T. conceived experimental designs and edited manuscript. D.M.H. provided PHF1 and MC1 antibodies, conceived experimental designs, and edited the manuscript. A.S.Y. supervised the project.

---

**Fig. S1.**
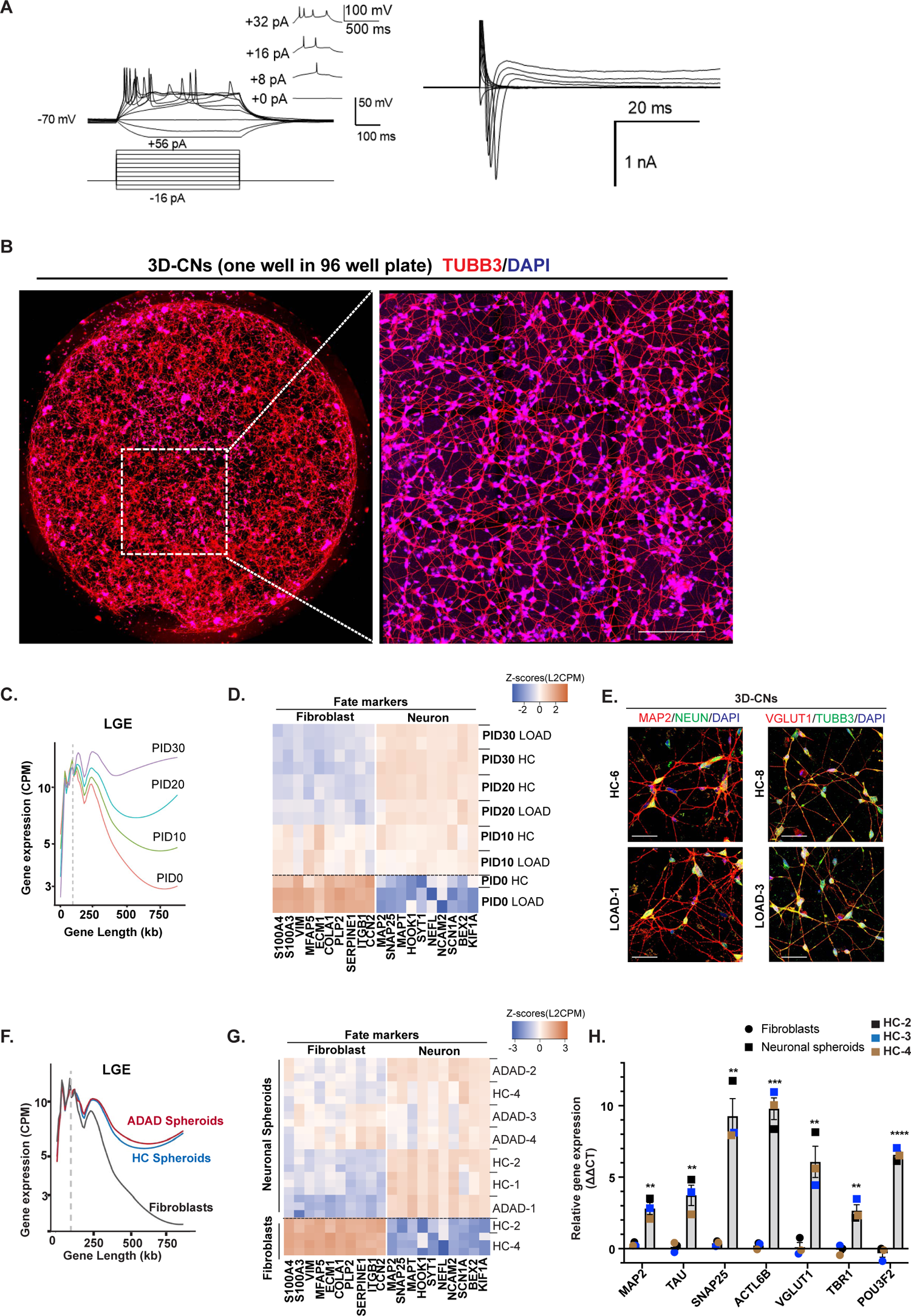
Direct reprogramming of patient fibroblasts into 3D-CNs and spheroids (related with Fig. 1) **A.** Left: Current-clamp whole cell recording from converted neurons (PID30) in 2D culture environment co-cultured with human astrocytes showing multiple action potentials (APs). Right: Voltage-clamp whole cell recording revealed the inward and outward currents. **B.** Left: Representative images of a thin gel culture (PID28) in one whole well of a 96-well plate. TUBB3 antibody was used to show the neuronal morphology. Right: A magnified image showing the boxed region on the left. Scale bar: 250 µm. **C.** A LONGO plot depicting increased long gene expression (LGE) during neuronal reprogramming in thin gel. **D.** Heatmap shows downregulation of fibroblast markers and upregulation of neuronal markers during neuronal reprogramming in thin gel. Replicates from an AD line and HC line were used for analysis in C and D. **E**. Immunostaining for MAP2, NEUN, VGLUT1 and TUBB3 in the thin gel culture of PID30 3D-CNs derived from LOAD patient fibroblasts. Scale bar: 50 µm. **F.** Expression of long genes in PID28 ADAD spheroids, PID28 HC spheroids, and their starting fibroblasts. **G.** A heatmap shows downregulation of fibroblast marker and upregulation of neuronal markers in PID28 neuronal spheroids compared to their starting fibroblasts. Spheroids derived from 4 independent ADAD patients and 3 HC individuals, and fibroblasts from 2 HC individuals were used for analysis in F & G. **H.** qPCR analysis of the mRNA expression of different neuronal markers in neuronal spheroids compared to their starting fibroblasts. n = 3 HC individuals. Mean ± SEM. **p < 0.01, ***p < 0.001 and ****p < 0.0001 by multiple unpaired t-tests.

**Fig. S2.**
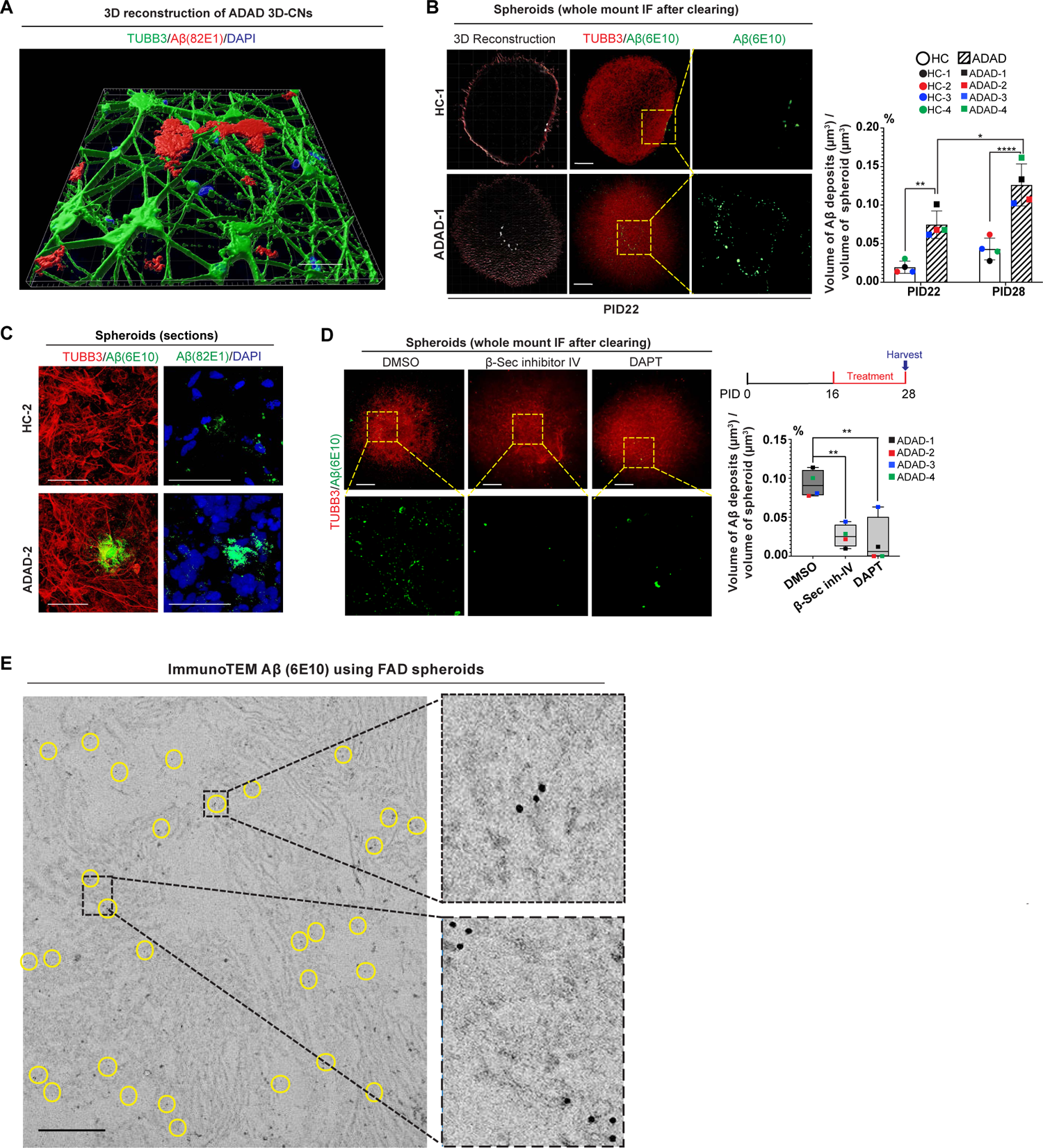
Detection of Aβ deposits in ADAD 3D-CNs and spheroids (related with Fig. 1) **A.** Tilted view of a 3D reconstruction image of ADAD 3D-CNs in Fig. 1G depicting the extracellular deposits labeled by 82E1 Aβ antibody (red). Scale bar: 50 µm. **B.** Representative images of whole mount immunostaining of Aβ followed by clearing of ADAD spheroids and sex- and age-matched HC spheroids at PID22. Scale bar: 500 µm. Representative images for Aβ deposition at PID28 can be found in Fig 1J. p-values were calculated by two-way ANOVA with Šídák’s multiple comparisons test. The adjusted p-value for comparison between PID22 HC and PID28 HC was not significant (p = 0.1855). *adjusted p < 0.05, **adjusted p < 0.01, and ****adjusted p < 0.0001. **C.** Representative immunofluorescence images of the sections of spheroids using two Aβ antibodies (left: 6E10; right: 82E1) showing the extracellular Aβ deposition. Scale bar: 50 µm. **D.** ADAD spheroids were treated with 5 µM β-secretase inhibitor IV or 40 µM DAPT starting at PID16 and immunostained for Aβ and TUBB3 at PID28. Adjusted p-values were calculated by one-way ANOVA with Dunnett’s multiple comparisons test. **p < 0.01. Scale bar: 500 µm. **E.** Transmission EM images from ADAD spheroids showing extracellular signals of immuno-gold-labeled Aβ antibody (6E10). Yellow circles highlight the nanogold particles conjugated to Aβ antibody. Scale bar: 1 µm. For quantification in B and D, n = 4 independent ADAD patients and 4 HC individuals.

**Fig. S3.**
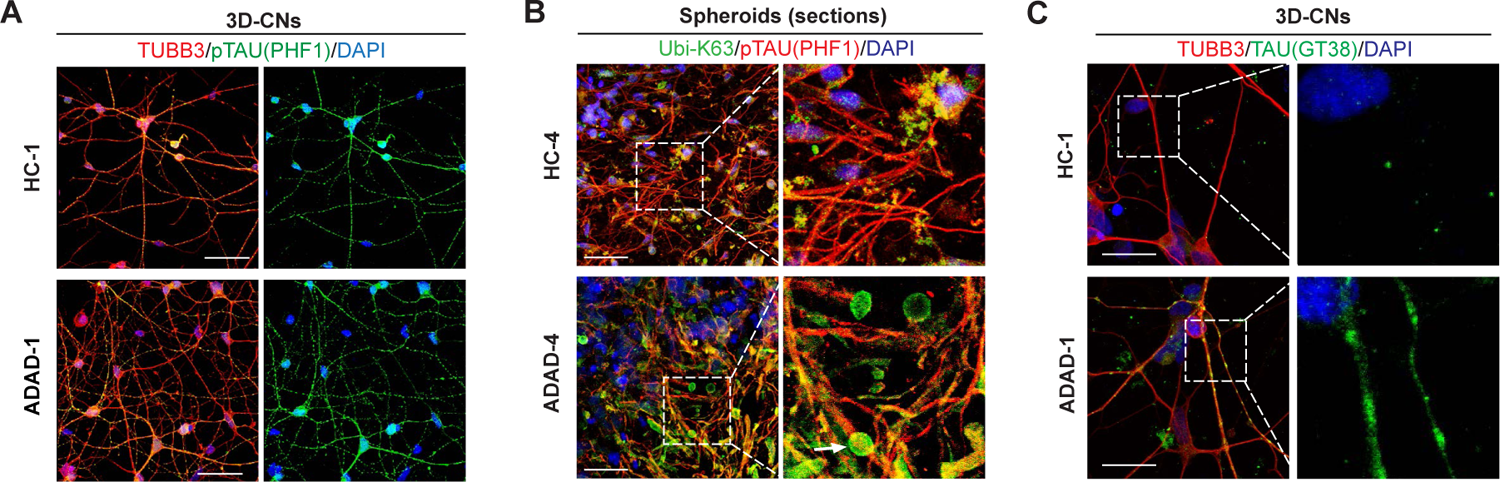
Identification of tauopathy in ADAD 3D-CNs and spheroids (related with Fig. 2) **A.** Representative immunostaining images of p-tau expression in thin gel using PHF1 p-tau antibody. Scale bar: 50 µm. **B.** Representative images of p-tau (PHF1) and K63-specific ubiquitin immunostaining in the sections of ADAD and HC spheroids. Scale bar: 50 µm. Arrow points to the enlarged tau blebs that are colocalized with K63-linked ubiquitin. **C.** Immunofluorescence staining images of AD-conformation-specific tau in ADAD and HC neurons using GT-38 tau antibody. Scale bar: 25 µm.

**Fig. S4.**
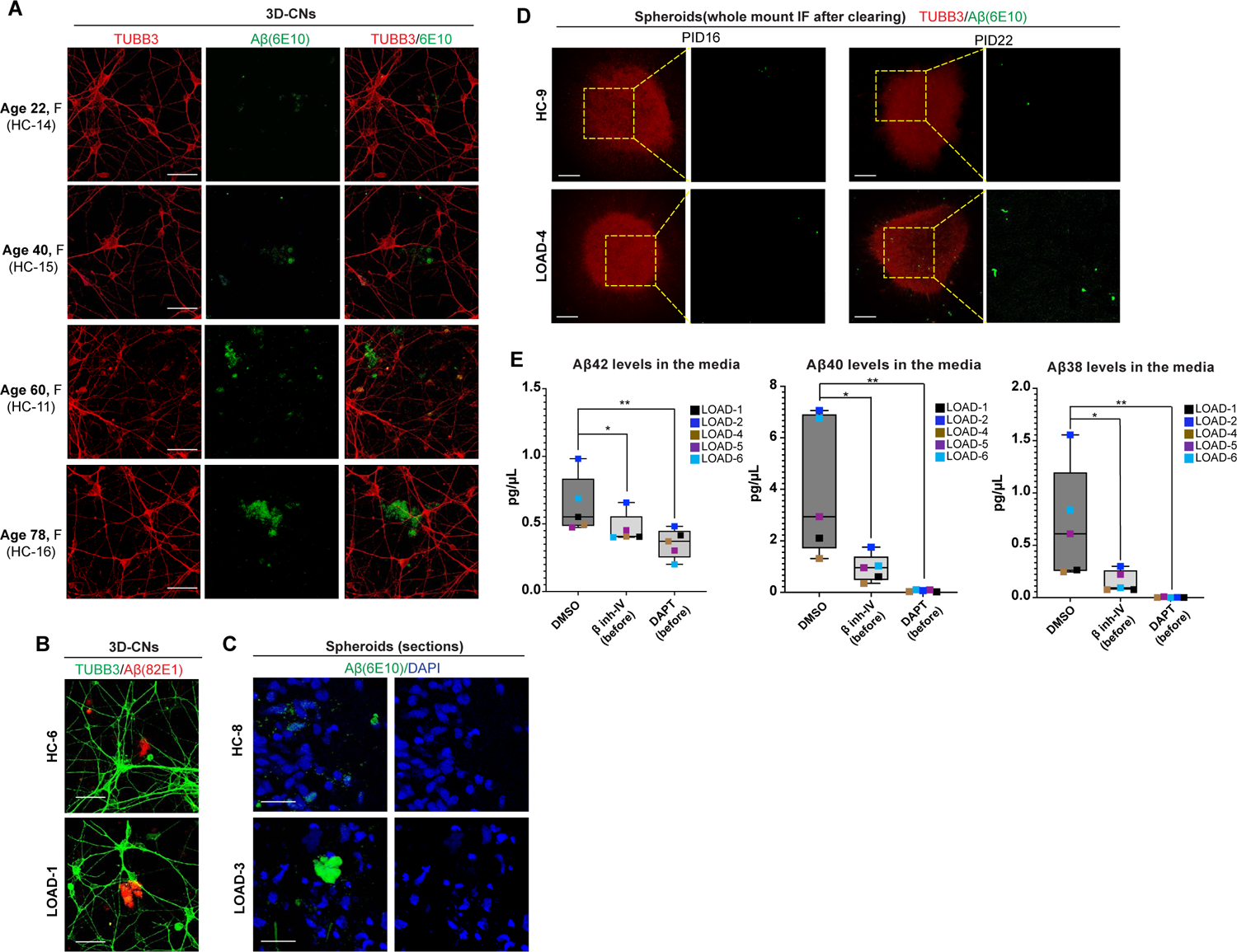
Detection of Aβ deposition in aging and LOAD 3D-CNs and spheroids (related with Fig. 4) **A.** Representative immunostaining images of Aβ deposition (6E10 antibody, green) in 3D-CNs derived from healthy individuals of different ages. F: female. Scale bar: 50 µm. **B.** Example of extracellular Aβ deposits labeled by 82E1 Aβ antibody in the thin gel culture of LOAD and HC at PID30. Scale bar: 50 µm. **C.** Representative immunostaining images of Aβ deposits on the sections of PID28 LOAD and HC spheroids using 6E10 Aβ antibody. Scale bar: 25 µm. **D.** Whole mount staining in PID16 and PID22 LOAD and HC spheroids using 6E10 Aβ antibody revealed negligible amount of Aβ deposition in both LOAD and HC group at PID16, but the amount became robust at PID22. Scale bar: 500 µm. **E.** Mass spectrometry quantification of Aβ42, Aβ40, and Aβ38 detected in the media of PID28 LOAD spheroids that were treated with β-secretase inhibitor IV or DAPT from PID16, corresponding to a time point before the onset of Aβ deposition. n = 5 independent LOAD patients. Adjusted p-values were calculated by one-way ANOVA with Dunnett’s multiple comparisons test. *p < 0.05 and **p < 0.01.

**Fig. S5.**
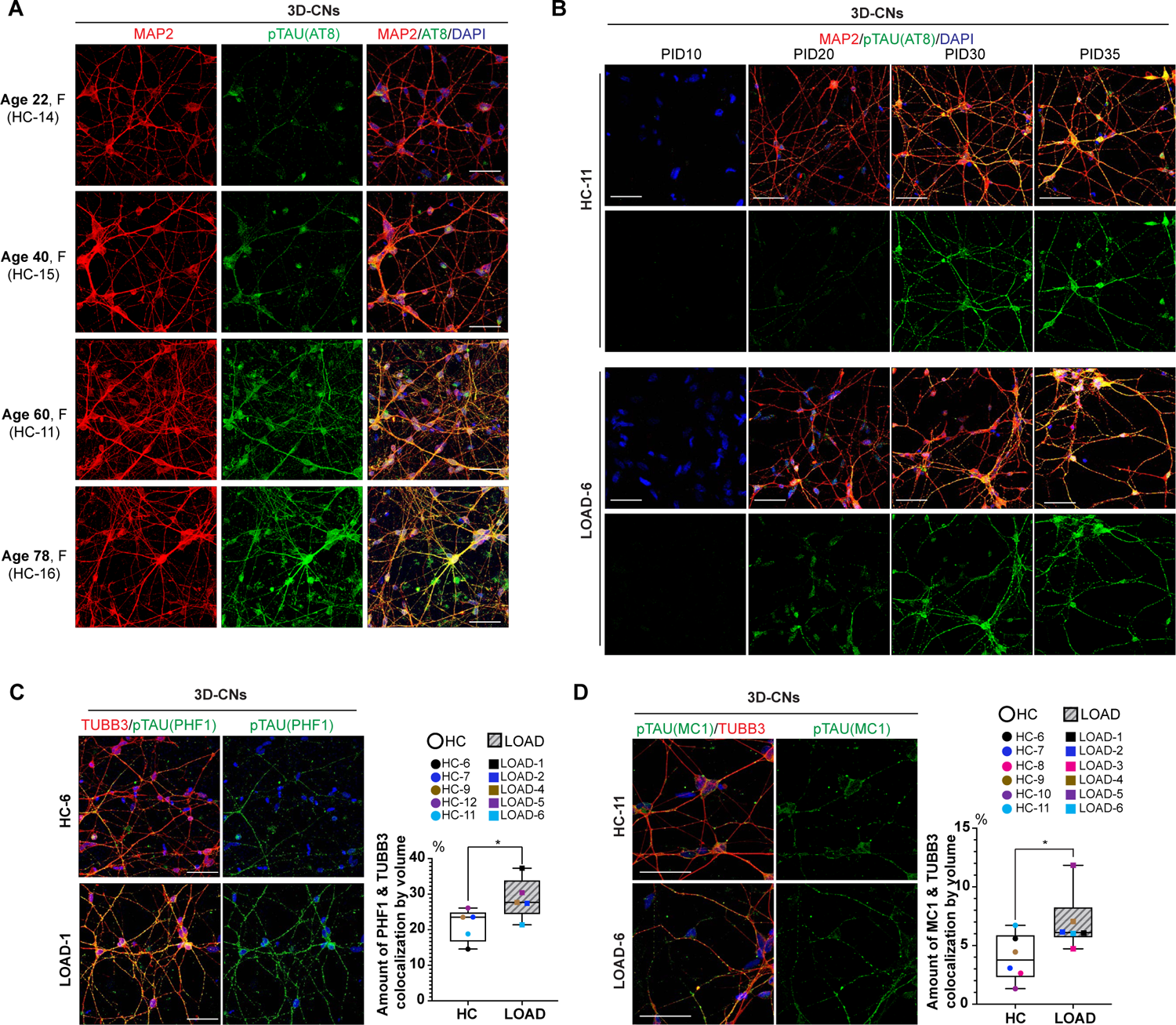
Tau signals in aging and LOAD 3D-CNs (related with Fig.4) **A.** Immunostaining of p-tau (AT8 antibody, green) in 3D-CNs reprogrammed from healthy individuals of advancing ages. F: female. Scale bar: 50 µm. **B.** p-Tau immunostaining using AT8 antibody at different PIDs of reprogramming showing p-tau signal at PID30 and PID35 in HC and LOAD 3D-CNs. Scale bar: 50 µm. **C.** Representative images of p-tau staining using PHF1 antibody in the PID30 LOAD and HC 3D-CNs. n = 5 independent LOAD patients and 5 HC individuals. *p < 0.05 by unpaired t-test. Scale bar: 50 µm. **D.** Misfolded tau was examined using MC1 tau antibody in both LOAD and HC 3D-CNs at PID30. n = 6 independent LOAD patients and 6 HC individuals. *p < 0.05 by unpaired t-test. Scale bar: 50 µm.

**Fig. S6.**
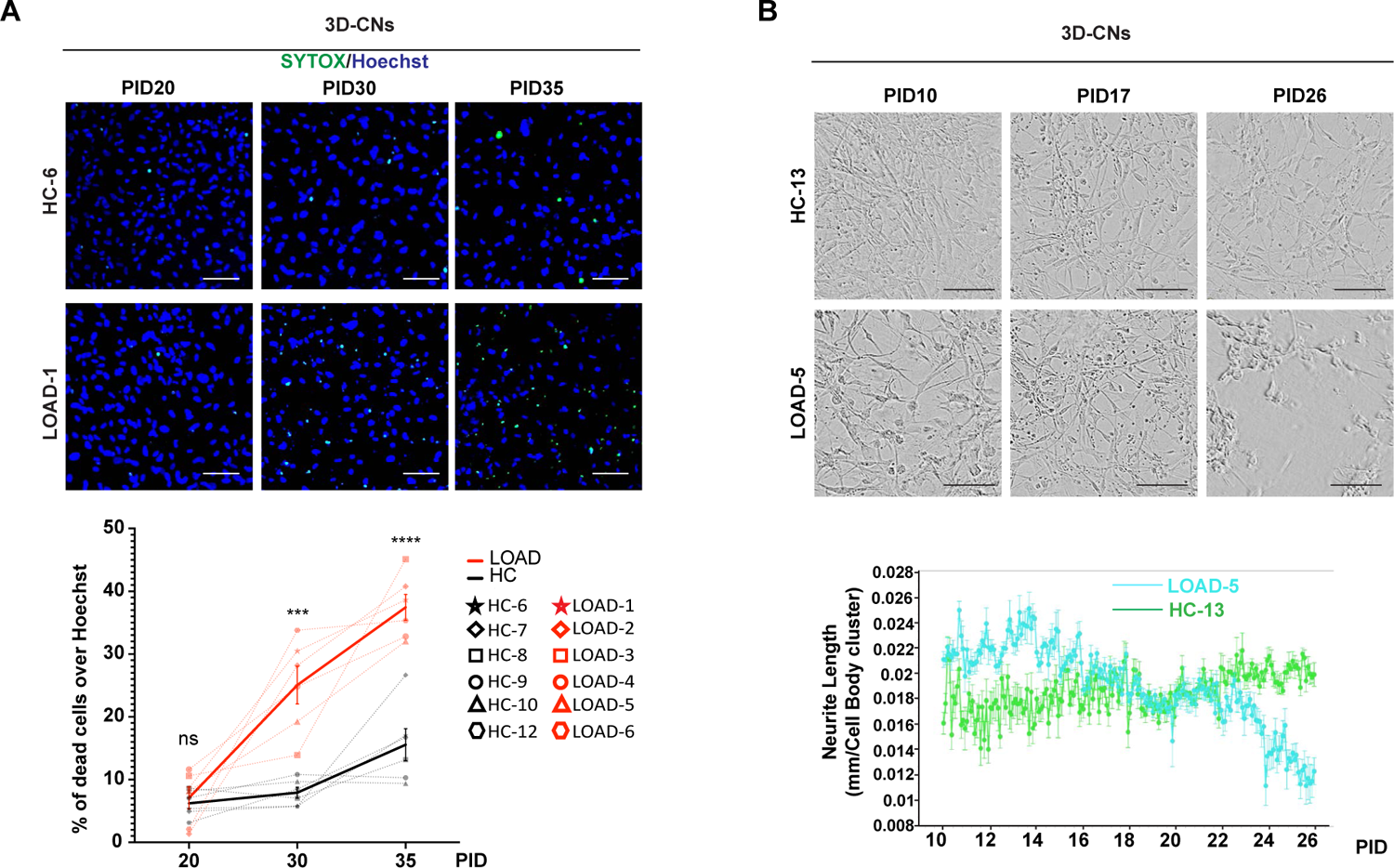
Spontaneous neuronal death in LOAD 3D-CNs (related with Fig. 4). **A.** Representative images of Sytox staining in LOAD and HC neurons reprogrammed in thin gel at different PIDs. Dead cells were labeled by Sytox-Green and nuclei were stained by Hoechst (blue). Statistics show the percentage of dead cells was significantly greater in LOAD 3D-CNs at PID30 and PID35 compared to the HC 3D-CNs at same PIDs. n= 6 independent LOAD patients and 6 HC individuals. Solid lines show the mean of each group while dotted lines represent each individual. Mean ± SEM. p-values were calculated by two-way ANOVA with Šídák’s multiple comparisons. ns: adjusted p > 0.05, ***adjusted p < 0.001, and ****adjusted p < 0.0001. Scale bar: 50 µm. **B.** Live cell imaging of LOAD and HC neurons at different PIDs during neuronal reprogramming. Quantification shows the decrease of neurite length in LOAD neurons starting from PID22. Normalized neurite extension was measured by the Neurotracker module of Incucyte Live-Cell analysis. Scale bar: 100 µm.

**Fig. S7.**
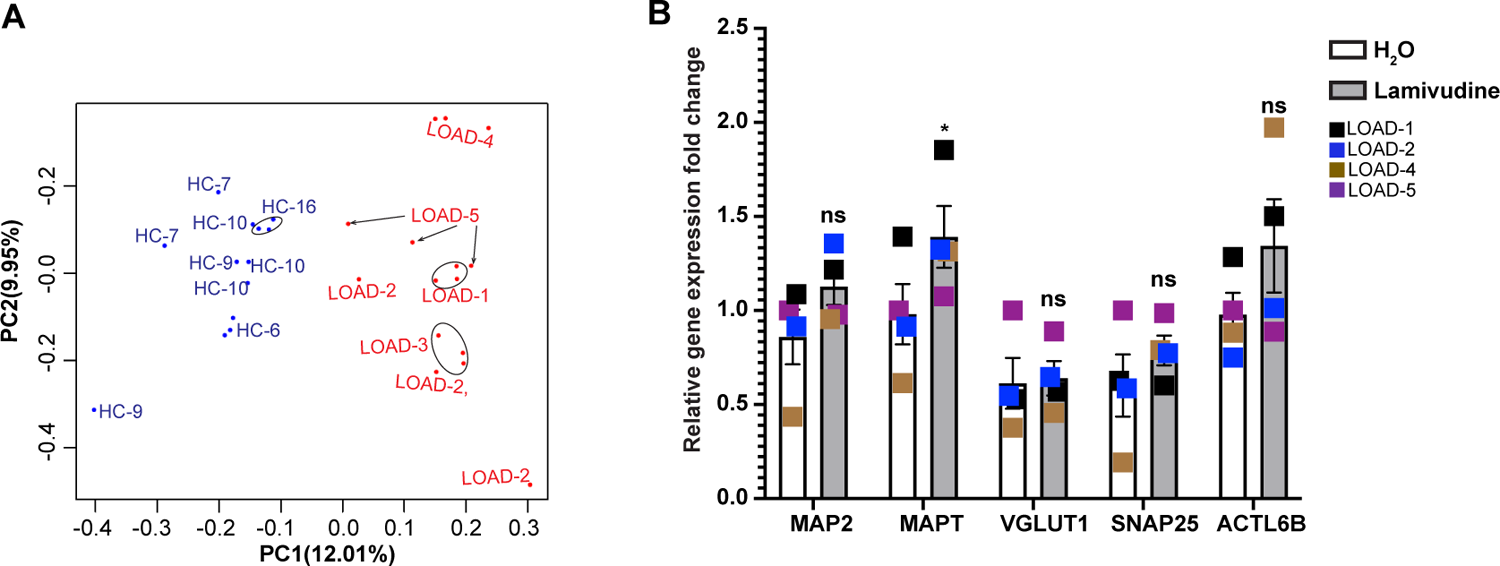
Lamivudine treatment does not dampen the neuronal identity in LOAD spheroids (related with Fig. 6). **A.** Principal component analysis of gene expression data of LOAD and HC spheroids. A total of 28 samples from 10 independent individuals (5 for LOAD and 5 for HC) were used for analysis. **B.** qPCR analysis of the mRNA expression of different neuronal markers in lamivudine treated spheroids compared to control spheroids that were treated with H_2_O. n = 4 LOAD patients. Mean ± SEM. ns (no significant difference) and *p < 0.05 by multiple unpaired t-tests.

**Table S1:** The list of AD and healthy control cell lines used in this study

| Sample ID | Fibroblast ID | Sex | Age at biopsy | Age at onset | Genotype | Origin |
| --- | --- | --- | --- | --- | --- | --- |
| ADAD-1 | AG06840 | male | 56 |  | ADAD, PSEN1 A246E | Coriell |
| HC-1 | AG04148 | male | 56 |  | Healthy Control | Coriell |
| ADAD-2 | UCL 33 | male | 58 | 49 | ADAD, APP V717I | UCL |
| HC-2 | AG04453 | male | 52 |  | Healthy Control, | Coriell BLSA |
| ADAD-3 | UCL 53 | male | 38 |  | ADAD, PSEN1 M146I | UCL |
| HC-3 | AG04060 | male | 44 |  | Healthy Control | Coriell |
| ADAD-4 | UCL 803 | female | 42 | 38 | ADAD, APP V717F | UCL |
| HC-4 | AG05838 | female | 36 |  | Healthy Control | Coriell |
| HC-5 | AG08260 | male | 61 |  | Healthy Control | Coriell |
| LOAD-1 | FA15-552 | male | 89 |  | Biomarker Positive, LOAD, APOE3/4 | Knight ADRC |
| HC-6 | FA18-634 | male | 89 |  | Biomarker Negative, Healthy, APOE3/3 | Knight ADRC |
| LOAD-2 | FA12-453 | female | 72 |  | Biomarker Positive, LOAD, APOE4/4 | Knight ADRC |
| HC-7 | AG13246 | female | 66 |  | Healthy Control | Coriell |
| LOAD-3 | AG05810 | female | 79 |  | LOAD, APOE3/4 | Coriell |
| HC-8 | AG11020 | female | 79 |  | Healthy Control | Coriell |
| LOAD-4 | FA17-623 | female | 87 |  | Biomarker Positive, LOAD, APOE3/3 | Knight ADRC |
| HC-9 | FA12-463 | female | 83 |  | Biomarker Negative, Healthy, APOE3/3 | Knight ADRC |
| LOAD-5 | FA18-633 | male | 70 |  | Biomarker Positive, LOAD | Knight ADRC |
| HC-10 | AG14251 | male | 68 |  | Healthy Control | Coriell |
| LOAD-6 | AG06869 | female | 60 |  | LOAD | Coriell |
| HC-11 | AG08379 | female | 60 |  | Healthy Control | Coriell |
| HC-12 | AG13369 | male | 68 |  | Healthy Control | Coriell |
| HC-13 | AG04356 | male | 74 |  | Healthy Control | Coriell |
| HC-14 | GM02171 | female | 22 |  | Healthy Control | Coriell |
| HC-15 | AG07307 | female | 40 |  | Healthy Control | Coriell |
| HC-16 | AG11246 | female | 78 |  | Healthy Control | Coriell |

Note: Biomarker positivity was defined based on amyloid imaging with PIB or AV45.

**Table S2:** The list of antibodies used in this study

| Antibody | Host | Supplier | Cat # | IF for 3D-CNs or<br>sections from<br>spheroid | Whole mount<br>IF staining for<br>spheroids |
| --- | --- | --- | --- | --- | --- |
| TUBB3 | Rabbit | Biologend | 802001 | 1:2500 | 1:1000 |
| TUBB3 | Mouse | Biologend | 801202 | 1:1000 | 1:500 |
| TUBB3 | Chicken | Novus | NB100-1612 | 1:1000 | 1:500 |
| NCAM1 | Mouse | Santa Cruz | SC-106 | 1:50 |  |
| VGLUT1 | Mouse | Millipore | MAB5502 | 1:200 |  |
| MAP2 | Rabbit | Cell signaling | #4542 | 1:200 |  |
| NeuN | Mouse | Chemicon | MAB377 | 1:200 |  |
| Amyloid-β (6E10) | Mouse | Biologend | 803001 | 1:200 | 1:100 |
| Amyloid-β (82E1) | Mouse | IBL | 10323 | 1:200 | 1:100 |
| P-TAU (AT8) | Mouse | Invitrogen | mn1020 | 1:200 |  |
| P-TAU (PHF1) | Mouse | Gift from Peter Davis | N/A | 1:400 |  |
| TAU (MC1) | Mouse | Gift from Peter Davis | N/A | 1:400 |  |
| K63-linked<br>UBIQUITIN | Rabbit | Abcam | ab179434 | 1:200 |  |
| TAU(GT38) | Mouse | Abcam | ab246808 | 1:200 |  |

**Table S3:**
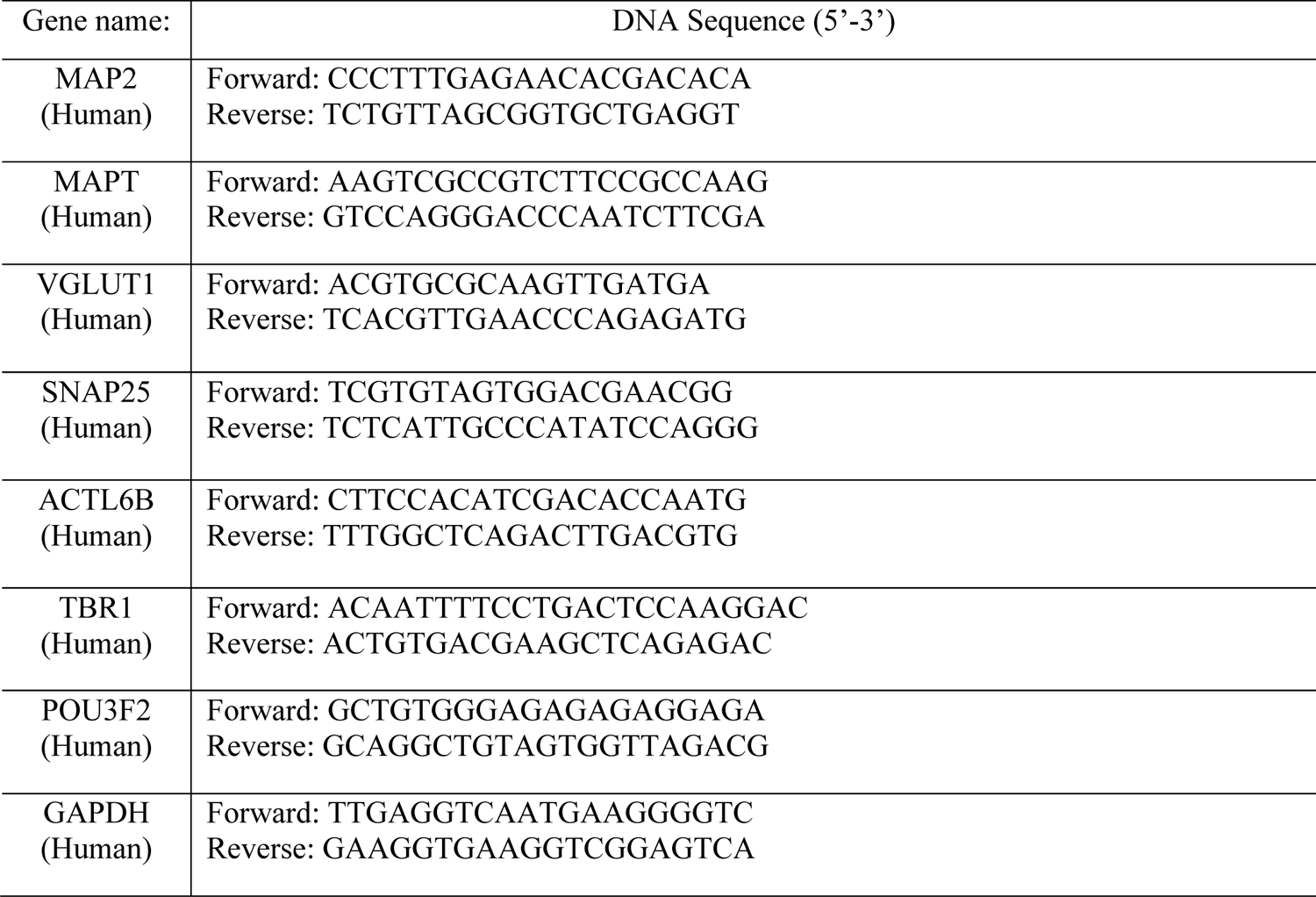
The list of qPCR primers used in this study

